# Synaptic and Gene Regulatory Mechanisms in Schizophrenia, Autism, and 22q11.2 CNV Mediated Risk for Neuropsychiatric Disorders

**DOI:** 10.1101/555490

**Authors:** Jennifer K. Forsyth, Daniel Nachun, Michael J. Gandal, Daniel H. Geschwind, Ariana E. Anderson, Giovanni Coppola, Carrie E. Bearden

**Affiliations:** Department of Psychiatry & Biobehavioral Sciences, University of California, Los Angeles, Los Angeles, CA 90095; Department of Human Genetics, University of California, Los Angeles, Los Angeles, CA 90095; Department of Neurology, University of California, Los Angeles, Los Angeles, CA 90095; Brain Research Institute, University of California, Los Angeles, Los Angeles, CA 90095; Department of Psychology, University of California, Los Angeles, Los Angeles, CA 90095

**Author notes:** Corresponding Authors: Jennifer K. Forsyth, Ph.D. and Carrie E. Bearden, Ph.D., 760 Westwood Plaza, Los Angeles, CA 90095, (J.K.F); (C.E.B).

**Keywords:** 22q11.2 copy number variants, schizophrenia, autism spectrum disorder, genetic risk, functional genomics, brain development

## Abstract

**Background:** 22q11.2 copy number variants (CNVs) are among the most highly penetrant genetic risk variants for developmental neuropsychiatric disorders such as schizophrenia (SCZ) and autism spectrum disorder (ASD). However, the specific mechanisms through which they confer risk remain unclear.

**Methods:** Using a functional genomics approach, we integrated transcriptomic data from the developing human brain, genome-wide association findings for SCZ and ASD, protein interaction data, and pathophysiological signatures of SCZ and ASD to: 1) organize genes into the developmental cellular and molecular systems within which they operate; 2) identify neurodevelopmental processes associated with polygenic risk for SCZ and ASD across the allelic frequency spectrum; and 3) elucidate pathways and individual genes through which 22q11.2 CNVs may confer risk for each disorder.

**Results:** Polygenic risk for SCZ and ASD converged on partially overlapping gene networks involved in synaptic function and transcriptional regulation, with ASD risk variants additionally enriched for networks involved in neuronal differentiation during fetal development. The 22q11.2 locus formed a large protein network that disproportionately affected SCZ- and ASD-associated neurodevelopmental networks, including loading highly onto synaptic and gene regulatory pathways. *SEPT5, PI4KA*, and *SNAP29* genes are candidate drivers of 22q11.2 synaptic pathology relevant to SCZ and ASD, and *DGCR8* and *HIRA* are candidate drivers of disease-relevant alterations in gene regulation.

**Conclusions:** The current approach provides a powerful framework to identify neurodevelopmental processes affected by diverse risk variants for SCZ and ASD, and elucidate the mechanisms through which highly penetrant multi-gene CNVs contribute to disease risk.

## Introduction

Copy number variants (CNVs) at chromosome 22q11.2 are among the genetic variants most strongly associated with developmental neuropsychiatric disorders. The hemizygous deletion, 22q11.2 deletion syndrome (22q11DS), typically spans a gene-rich ~2.5 Mb region(1) and is the greatest known single genetic risk factor for schizophrenia (SCZ). Approximately 25% of 22q11DS patients develop a psychotic disorder by adulthood and ~0.5-1 % of patients with SCZ have 22q11DS(2). Interestingly, the reciprocal duplication (22q11DupS) may protect against SCZ(3,4), whereas both 22q11DS and 22q11DupS confer risk for autism spectrum disorder (ASD;5). As psychiatry increasingly looks towards precision medicine approaches to therapeutic development, clarifying the mechanisms through which 22q11.2 CNVs confer risk for SCZ and ASD, and how these mechanisms converge with risk derived from broader variants associated with each disorder, is a critical goal.

Our knowledge of the genetic architectures of both SCZ and ASD has advanced considerably in the last decade. It is now clear that SCZ and ASD are both highly polygenic and genetically heterogeneous disorders, with risk variants for each disorder spanning common (i.e., present in >1% of the population), rare (i.e., present in <1% of the population), and de novo variants (i.e., new mutations in offspring), in addition to CNVs(6). Despite this polygenicity, risk variants for SCZ and ASD each appear to converge on a subset of biological pathways. Neuronal and synaptic genes have been repeatedly implicated in SCZ, including the N-methyl-D-aspartate receptor (NMDAR) and activity-regulated cytoskeleton-associated protein (Arc) complexes, and targets of the fragile X mental retardation protein (FMRP;4,7–11). Similarly, neuronal and synaptic genes have been implicated in ASD, along with chromatin and transcription regulators, and FMRP targets(5,12–16). However, many genes in these gene-sets are developmentally regulated, and SCZ and ASD are both neurodevelopmental in origin. Indeed, brain development is tightly controlled through changes in transcription factor activity and chromatin condensation that alter gene expression patterns at precise times in development(17). These changes program the progression of stem cell proliferation, neuronal differentiation, and synaptogenesis during prenatal development, and myelination and synaptic refinement during postnatal development. Examining the convergence of SCZ and ASD risk variants on groups of developmentally co-expressed genes may therefore yield fundamental insights into the neurodevelopmental pathogenesis of each disorder (e.g.,15,16), and provide a powerful lens to understand how 22q11.2 CNVs confer risk for each disorder.

Here, we used network analyses of transcriptomic data from the developing human brain(17) to organize genes into the developmental cellular and molecular systems within which they operate(18). We then conducted a comprehensive examination of neurodevelopmental processes associated with risk variants for SCZ and ASD across the allelic frequency spectrum. Finally, we identified the brain developmental 22q11.2 protein network, and developed a framework to leverage this disease-informed lens to predict pathways and individual genes underlying 22q11.2 CNV-mediated risk for SCZ and ASD.

## Methods and Materials

### Defining Neurodevelopmental Co-Expression Networks

Networks of developmentally co-expressed genes were constructed using weighted gene co-expression network analysis (WGCNA) of BrainSpan data, which is the largest, publicly-available developmental brain transcriptome dataset derived from healthy human donors(17). Samples spanning the full period of brain development (i.e., ~6 post-conception weeks-30 years; 1061 samples) were included in the analysis. Briefly, correlation coefficients were computed between expression levels for each gene, and connectivity measures were calculated by summing the connection strength of each gene with every other gene. Genes were clustered based on connectivity to identify groups of co-expressed genes (i.e., modules), and the mean expression of the first principal component of each module (i.e., module eigengene) per developmental stage was used to summarize the developmental trajectory of each module. Genes in each module were annotated for enrichment of: 1) brain cell-type markers(19); 2) gene ontology (GO) using g:Profiler(20); and 3) genes expressed in specific human brain regions and developmental periods using the Specific Expression Analysis tool(21). See Supplementary Methods for details.

### Identifying Neurodevelopmental Networks Associated with SCZ and ASD

Summary statistics, gene-lists, and/or variant lists from genome-wide association studies of SCZ and ASD were compiled and tested for module enrichment using methods appropriate to each data type. Briefly, when summary level statistics for common(8,11,22) and rare variants(23) were available, they were tested for enrichment using MAGMA(24). De novo mutations (DNMs) were compiled for SCZ(9,25–32) and ASD(33,34); the affected gene and predicted consequence of each DNM were annotated using Ensembl Variant Effect Predictor (VEP) and tested for enrichment using the DenovolyzeR package(35). Genes in SCZ-(4) and ASD-associated(14) CNV loci were tested for enrichment using logistic regression, controlling for gene length. Genes containing more deleterious ultra-rare variants in SCZ patients than controls(10); manually curated high-confidence genes implicated in ASD according to the SFARI gene database(36); and genes associated with ASD across rare inherited, de novo, and copy number variants(14) were formulated as hypergeometric distributions and tested for module enrichment using Fisher’s exact tests with the GeneOverlap package(37). See Table 1 for summary of data sources and Supplementary Methods for details.

**Table 1.** Genome-wide SCZ and ASD Risk Variant Datasets Tested for BrainSpan Module Enrichment.

| Data | Reference | Variant Type | Description | Patient N | Control N | Analysis |
| --- | --- | --- | --- | --- | --- | --- |
| SCZ |  |  |  |  |  |  |
| SCZ PGC-deCODE* | 11 | Common | PGC + deCODE meta-analysis | 36,989 | 113,075 | MAGMA |
| SCZ CLOZUK-PGC* | 8 | Common | CLOZUK + PGC meta-analysis | 40,675 | 64,643 | MAGMA |
| SCZ Richards | 23 | Rare | Rare variants (MAF<1%) | 5,585 | 8,103 | MAGMA |
| SCZ Genovese | 10 | Ultra-rare | 43 genes enriched for deleterious ultra-rare variants (p<0.01) | 4,877 | 6,242 | Hypergeometric Test |
| SCZ DNM | 9, 25-32 | De novo | Compilation of DNMs | 1,139 | NA | DenovolyzeR |
| SCZ CNV | 4 | CNV | 13 unique loci (FDR<0.05) | 21,094 | 20,227 | Logistic Regression |
| ASD |  |  |  |  |  |  |
| ASD iPSYCH-PGC | 22 | Common | iPSYCH + PGC meta-analysis | 18,381 | 27,969 | MAGMA |
| ASD SFARI | www.sfari.org | Gene list | 80 literature-curated genes (score 1 or 2) | NA | NA | Hypergeometric Test |
| ASD Sanders | 14 | Rare | 65 genes (FDR<0.1) | 3,982 | 1,911 | Hypergeometric Test |
| ASD DNM | 33, 34 | De novo | Compilation of DNMs | 4,424 | NA | DenovolyzeR |
| ASD CNV | 14 | CNV | 9 unique loci, +3 nested loci, in SSC and AGP cohorts (FDR<0.05) | 4,687 | 2,100 | Logistic Regression |

\*The PGC-deCODE meta-analysis includes 1,513 SCZ cases and 36,989 controls from the deCODE study not in the CLOZUK-PGC study and the SCZ CLOZUK-PGC meta-analysis includes 5,220 SCZ cases and 18,823 controls not in the SCZ PGC-deCODE study.

### Characterizing the 22q11.2 Locus

To examine whether any BrainSpan module was over-represented among the 22q11.2 genes, given the number of genes in each module, or whether SCZ- and ASD-associated modules were over-represented overall, logistic regression controlling for gene length was conducted for each module, and for all disease-associated modules together, respectively.

To characterize the total protein-coding gene content of the 22q11.2 locus and its gene content per module relative to all regions of the genome, the 22q11.2 locus was compared to a distribution generated by simulating CNVs of the same size (~2.57 mb) in 250 kb sliding steps across the genome (i.e., 12,149 genome-wide regions), using the GenomicRanges package(38). The percentile ranks of the 22q11.2 locus overall, and per module, were calculated relative to this distribution. A Mann-Whitney U test examined whether the 22q11.2 locus percentile ranks were higher for SCZ- and ASD-associated modules, overall, than for non-associated modules.

### Characterizing the 22q11.2 Brain Development PPI Network

Genes form protein-protein interaction (PPI) networks that fulfill critical biological roles, and perturbing protein networks is a common characteristic of a majority of disease-associated mutations(39). However, protein interactions are dependent on the tissue and temporal availability of each protein(40). We therefore identified the brain development protein-protein interaction network (BD-PPI network) for the 22q11.2 locus (i.e., 22q_BD-PPI_ network) and for: 1) all ~2.57 Mb regions of the genome, as described above (i.e., Genomic-Region_BD-PPI_ networks); and 2) 10,000 random gene-sets, equal in number of seed BrainSpan genes to the 22q11.2 locus (i.e., Gene-Set_BD-PPI_ networks).

The 22q_BD-PPI_ network was first tested for over-representation in any BrainSpan modules, given the number of genes in each module, using hypergeometric overlap tests. To characterize the 22q11.2 region relative to all regions of the genome, and more specifically, the connectivity of 22q11.2 genes within protein networks relative to all BrainSpan genes, we then calculated the 22q_BD-PPI_ network percentile rank per module relative to the Genomic-Region_BD-PPI_ and Gene-Set_BD-PPI_ networks, respectively. Mann-Whitney U tests examined whether the 22q_BD-PPI_ network percentile ranks were higher overall for SCZ- and ASD-associated modules than for non-associated modules.

To generate BD-PPI networks, protein pairs were compiled from the BioGRID 3.4(41) and InWeb3 PPI databases(42). PPI pairs were thresholded by BrainSpan-derived brain development co-expression pairs (Spearman’s p>0.7;16) to identify high-confidence PPI pairs likely to exist during brain development. BD-PPI networks were identified by extracting the seed genes and corresponding BD-PPI pairs of each genomic region or gene-set. See Supplementary Methods for details.

### Prioritizing Disease-Relevant 22q11.2 Genes

To prioritize individual 22q11.2 genes likely to drive risk for SCZ and ASD, the number of genes per BrainSpan module in each individual 22q11.2 gene’s BD-PPI network was compared to that for every other BrainSpan gene. BD-PPI networks of genes above the 95th percentile for SCZ- or ASD-associated modules were examined for GO term enrichment using g:Profiler.

### Integrating SCZ and ASD Pathophysiological Signatures

Significantly down- and up-regulated genes in SCZ and ASD post-mortem cortex were obtained from (43) and first tested for BrainSpan module enrichment using hypergeometric overlap tests.

The 22q_BD-PPI_ network overlapping down-regulated genes in SCZ and ASD was then characterized relative to the above described Genomic-Region_BD-PPI_ and Gene-Set_BD-PPI_ networks. Similarly, the number of genes in each individual 22q11.2 gene’s BD-PPI network overlapping down-regulated genes in SCZ and ASD was compared to that for every other BrainSpan gene.

## Results

### Genetic Risk for SCZ and ASD Converge on Partially Overlapping Neurodevelopmental Processes

Eighteen BrainSpan modules were identified across 17,216 genes (Fig. S1A; Table S1; 4 genes were not assigned to any module) with distinct developmental expression trajectories, cell-type enrichment patterns (Fig. S1B), and functional characteristics (Table S2), capturing broad aspects of neurotypical development.

SCZ and ASD risk variant enrichment for BrainSpan modules is summarized in Fig. 1. SCZ-associated variants largely converged on 3 modules. SCZ-associated common variants and CNVs were significantly over-represented in M15 (FDR corrected <0.05 for number of modules), a neuronal marker-enriched module that increased in expression through adolescence and was enriched for genes expressed during postnatal development (Figs. 2A-C, S2M-N). GO analysis indicated that M15 was involved in synaptic transmission, modulation of neurotransmission, and learning and memory. SCZ common variants were additionally over-represented in M13, a neuronal module that increased in expression during prenatal development, with relatively stable postnatal expression (Fig. 2D-F). Consistent with this, M13 was enriched for genes expressed during late fetal and postnatal development (Fig. S2K). M13 was enriched for synaptic transmission and regulation of the membrane potential, and its hub genes included *ATP1B1* and *KCNMA1*, which encode components of the Na+/K+ pump and a voltage-gated K+ channel, respectively, that critically regulate the electrochemical gradient(44,45). Thus, M13 appears to be involved in regulating basic neuronal excitability. Finally, SCZ common variants were significantly over-represented in M7, with rare variants showing nominal enrichment. M7 was involved in RNA processing and binding, and its hub genes included *POLR2B*, which encodes a major subunit of RNA polymerase II, the central enzyme responsible for transcribing mRNA from DNA, and *CREB1*, which encodes the transcription factor CREB, known to regulate neuronal differentiation, survival, and plasticity(46; Fig. 2G-I). M7 showed its highest expression during fetal development but was minimally enriched for genes associated with any specific period of development (Fig. S2G). Together, this suggests that M7 regulates gene expression and RNA processing across development.

**Figure 1.**
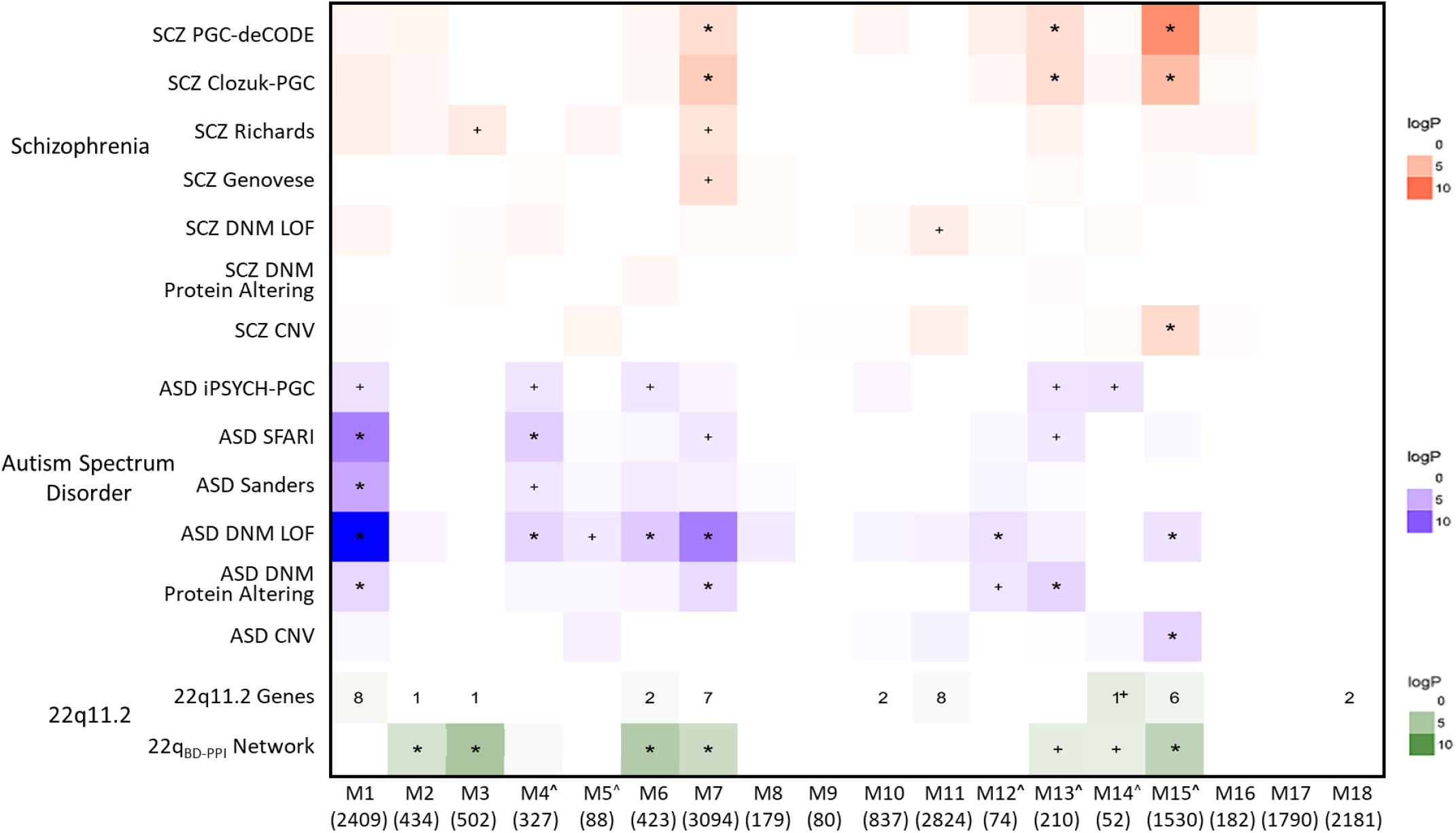
Genetic Risk for SCZ and ASD Converge on Partially Overlapping and Partially Distinct Neurodevelopmental Networks. BrainSpan module enrichment for SCZ (red) and ASD (blue) risk variant datasets, 22q11.2 genes, and the 22q11.2 brain-development PPI network (22q_BD-PPI_ network; green). Log p values are indicated by color for all datasets with enrichment odds ratios >1.0. To show potential convergence of enrichment across datasets, all enrichment values for p<0.05 are annotated (+p<0.05; *FDR corrected p<0.05 for number of modules). Numerical values denote number of 22q11.2 genes in each module. ^Module enriched for neuronal cell-type markers.

Risk variants for ASD showed overlapping enrichment for SCZ-associated modules, as well as enrichment for 4 distinct modules. Thus, loss-of-function (LoF) DNMs in ASD were over-represented in M7, with protein-altering DNMs, more broadly, and SFARI-defined ASD genes also nominally enriched in M7. Protein-altering DNMs in ASD were over-represented in M13, with common variants and SFARI-defined ASD genes nominally enriched. LoF DNMs and ASD-associated CNVs (many of which are also associated with SCZ; Table S3) were also over-represented in M15.

The 4 additional modules significantly associated with ASD were M1, M4, M6, and M12. Thus, rare variants, broadly, and LoF and protein-altering DNMs in ASD were consistently over-represented in M1, with common variants also nominally enriched. M1 peaked in expression during early-mid fetal development and was enriched for genes showing fetal-specific expression (Fig. S2A). M1 was involved in transcription, chromatin modification, and gene expression, and its hub genes included the transcription factors *SOX11* and *SOX4*, which are critical for promoting neuronal differentiation(47; Fig. 2J-L). Further, M1 contained numerous genes encoding subunits of the BAF chromatin remodeling complexes, which program gene expression changes that mediate embryonic stem cell differentiation into neurons(48). Thus, M1 appears to be particularly important for regulating fetal gene expression patterns that direct neuronal differentiation. LoF DNMs and SFARI-defined ASD genes were additionally over-represented in M4, with common and rare variants also nominally enriched. M4 was enriched for neuronal markers, peaked in expression during mid-fetal development, and was involved in neuronal differentiation and neuron projection development (Fig. 2M-O). Consistent with this, M4 was enriched for genes with fetal-specific expression (Fig. S2C), and its hub genes included DOK4, ARHGAP33, and TRIO, which are involved in axon and dendrite morphogenesis(49–51). LoF DNMs in ASD were also over-represented in M6, with common variants nominally enriched. M6 showed maximal expression during fetal development but was not enriched for genes expressed in any specific period of development (Fig. S2E), and was involved in translation and protein catabolism (Fig. 2P-R). Finally, LoF DNMs in ASD were also over-represented in M12, with protein-altering DNMs, more broadly, nominally enriched. M12, a neuronal module that is closely related to M13 (Fig. S1A,B), showed an increasing trajectory of expression until largely plateauing in late fetal development, and was involved in regulating the membrane potential (Fig. 2S-U). Module enrichment for gene-sets implicated in prior genetic studies of SCZ or ASD were consistent with GO analyses (Fig. S1C).

**Figure 2.**
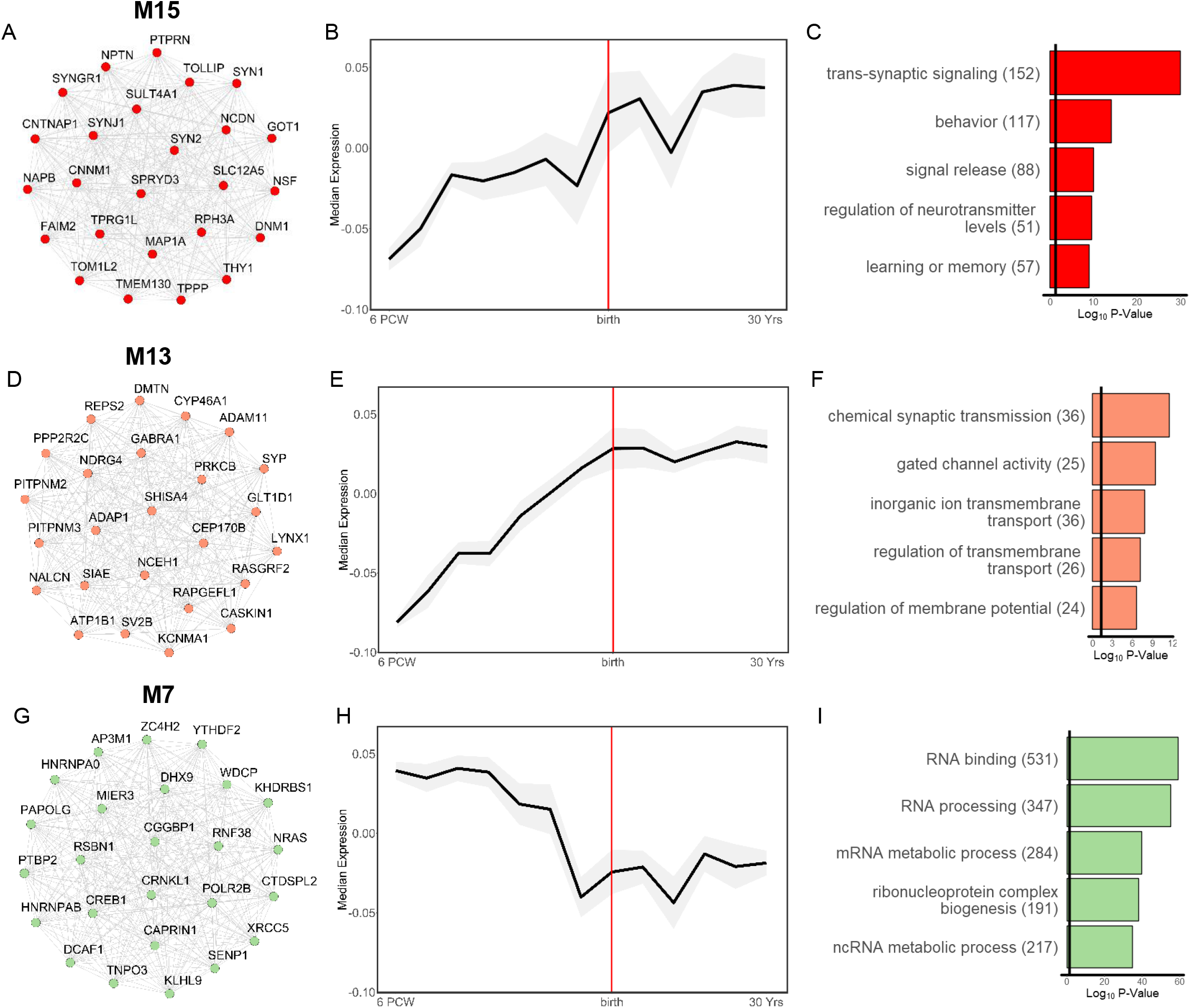
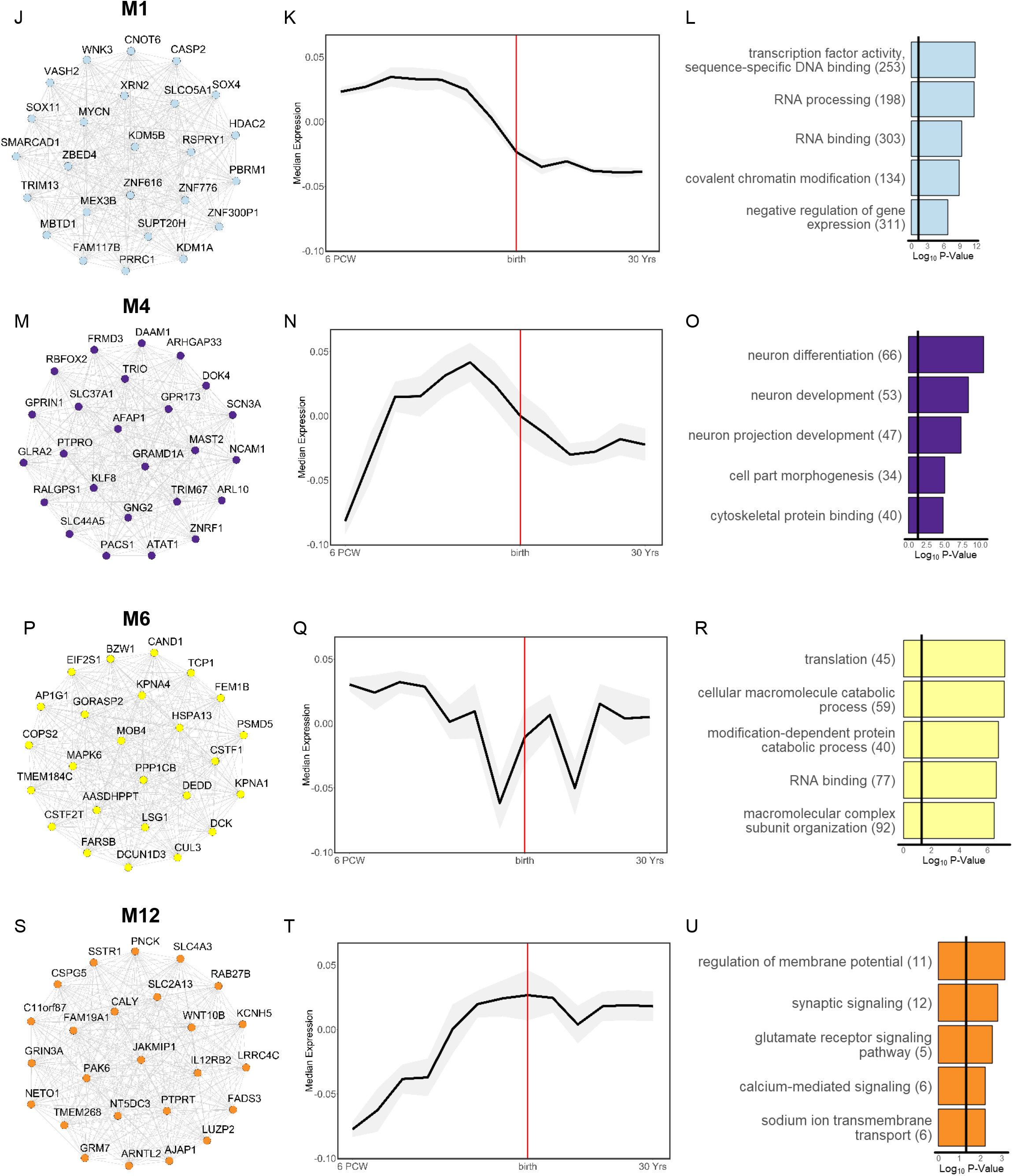
Characterization of SCZ- and ASD-Associated BrainSpan Modules. Left: top 25 hub genes, Middle: developmental expression trajectory (mean ± standard deviation in grey shading), and Right: top 5 significantly enriched biological process and molecular function gene ontology (GO) terms for key SCZ-associated (M15, M13, M7) and ASD-associated (M15, M13, M7, M1, M4, M6, M12) BrainSpan modules. Hub genes and GO terms are colored by module. Edges between hub genes denote positive correlations between genes.

As expected, LoF and protein-altering DNMs in control subjects were not over-represented in any SCZ- or ASD-associated modules, and no modules were significantly enriched for synonymous DNMs in ASD, SCZ, or control subjects, *p*s>0.05.

Thus, polygenic risk for SCZ and ASD appear to both coalesce on neurodevelopmental networks involved in postnatal neuronal and synaptic function and pan-developmental regulation of gene expression; however, risk variants for ASD additionally converge on networks involved in early neuronal differentiation and synapse formation.

### The 22q_BD-PPI_ Network Loads Highly onto SCZ- and ASD-Associated Modules

Forty-six protein-coding genes are contained in the ~2.57 Mb 22q11.2 locus; relative to all equal-sized regions of the genome, this was in the 93.58 percentile. Thirty-eight 22q11.2 genes were indexed in BrainSpan (Table 2). There were nominally more 22q11.2 genes than expected in M14, given the number of genes in each module (i.e., 1 gene; SEPT5; *p*=0.02; Fig. 1), but this did not survive FDR correction. Similarly, although 23 22q11.2 genes were in SCZ- and/or ASD-associated modules, this was not significantly more than expected, *p*>0.05. Relative to all equal-sized regions of the genome (~2.57 Mb), 22q11.2 gene content was above the 90^th^ percentile for 6 BrainSpan modules, including 4 modules associated with SCZ or ASD (M1, M6, M7, M15; Table S4). However, the genomic-region rank percentiles of the 22q11.2 locus for SCZ and ASD-associated modules, overall, were not significantly higher than for non-disease associated modules, *p*>0.05.

**Table 2.** BrainSpan Module Membership of 22q11.2 Genes. The 38 22q11.2 genes indexed in BrainSpan are annotated by module assignment, kME score (i.e., correlation with parent module eigengene) and pLI score (i.e., loss-of-function intolerance; bolded text indicates predicted pLI gene). ^Module enriched for neuronal cell-type markers.

| Module | Gene | Definition | kME Score | pLI Score |
| --- | --- | --- | --- | --- |
| M1 | ZNF74 | zinc finger protein 74 | 0.843 | 0.0004 |
|  | DGCR8 | DGCR8, microprocessor complex subunit | 0.757 | **0.9999** |
|  | SLC25A1 | solute carrier family 25 member 1 | 0.754 | 0.6335 |
|  | ARVCF | armadillo repeat gene deleted in velocardiofacial syndrome | 0.669 | 0.0000 |
|  | CLTCL1 | clathrin heavy chain like 1 | 0.607 | 0.0000 |
|  | KLHL22 | kelch like family member 22 | 0.562 | 0.7026 |
|  | TMEM191A | transmembrane protein 191A | 0.182 | NA |
|  | SCARF2 | scavenger receptor class F member 2 | 0.009 | NA |
| M2 | CRKL | v-crk avian sarcoma virus CT10 oncogene homolog-like | 0.511 | 0.1635 |
| M3 | CDC45 | cell division cycle 45 | 0.775 | 0.0691 |
| M6 | MRPL40 | mitochondrial ribosomal protein L40 | 0.779 | 0.0201 |
|  | ESS2 | DiGeorge syndrome critical region gene 14 | 0.752 | 0.0000 |
| M7 | UFD1 | ubiquitin recognition factor in ER associated degradation 1 | 0.797 | **0.9974** |
|  | SNAP29 | synaptosome associated protein 29 | 0.749 | 0.1517 |
|  | HIRA | histone cell cycle regulator | 0.635 | **1.0000** |
|  | RTL10 | retrotransposon Gag like 10 | 0.547 | 0.0000 |
|  | TRMT2A | tRNA methyltransferase 2 homolog A | 0.348 | 0.0000 |
|  | THAP7 | THAP domain containing 7 | 0.208 | 0.0293 |
|  | DGCR2 | DiGeorge syndrome critical region gene 2 | 0.165 | 0.0011 |
| M10 | AIFM3 | apoptosis-inducing factor, mitochondrion-associated, 3 | 0.941 | 0.0000 |
|  | P2RX6 | purinergic receptor P2X, ligand-gated ion channel, 6 | 0.799 | 0.0000 |
| M11 | PRODH | proline dehydrogenase (oxidase) 1 | 0.855 | 0.0000 |
|  | ZDHHC8 | zinc finger, DHHC-type containing 8 | 0.755 | **0.9890** |
|  | COMT | catechol-O-methyltransferase | 0.632 | 0.0008 |
|  | TXNRD2 | thioredoxin reductase 2 | 0.590 | 0.0000 |
|  | SERPIND1 | serpin peptidase inhibitor, clade D (heparin cofactor), member 1 | 0.416 | 0.0000 |
|  | C22orf39 | chromosome 22 open reading frame 39 | 0.243 | 0.0000 |
|  | DGCR6L | DiGeorge syndrome critical region gene 6-like | 0.172 | 0.1286 |
|  | GSC2 | goosecoid homeobox 2 | 0.146 | 0.0372 |
| M14^ | SEPT5 | septin 5 | 0.798 | **0.9146** |
| M15^ | PI4KA | phosphatidylinositol 4-kinase alpha | 0.881 | 0.0008 |
|  | TANGO2 | transport and golgi organization 2 homolog | 0.820 | 0.0000 |
|  | SLC7A4 | solute carrier family 7 member 4 | 0.722 | 0.0001 |
|  | RTN4R | reticulon 4 receptor | 0.573 | 0.5264 |
|  | MED15 | mediator complex subunit 15 | 0.381 | **0.9610** |
|  | LZTR1 | leucine zipper like transcription regulator 1 | 0.247 | 0.0000 |
| M18 | CLDN5 | claudin 5 | 0.434 | 0.7405 |
|  | TSSK2 | testis-specific serine kinase 2 | 0.382 | 0.0042 |

We next sought to characterize the 22q11.2 brain development PPI network (22q_BD-PPI_ network), given that perturbation of protein networks is a common characteristic of mutations associated with many diseases(39). The 22q11.2 locus formed a large BD-PPI network, consisting of 239 genes from 38 seed genes. Given the number of genes in each BrainSpan module, the 22q_BD-PPI_ network was significantly over-represented in multiple SCZ- and/or ASD-associated modules (M6, M7, M15), and nominally over-represented in M13 (Fig. 1). The 22q_BD-PPI_ network was also significantly over-represented in M2 and M3, and nominally over-represented in M14. However, some BrainSpan modules consist of genes that are more highly connected than other modules (i.e., have higher mean BD-PPI partners per gene, ANOVA *p*<2.0×10^-16^; Table S5), and highly connected genes are more likely to be found in the BD-PPI network of any random set of genes. This may be biologically meaningful, with gene networks that are more highly connected during brain development potentially more susceptible to perturbation. Nevertheless, to better understand the 22q_BD-PPI_ network relative to the rest of the genome, we therefore asked 2 additional questions: 1) how highly does the 22q_BD-PPI_ network load onto disease-associated modules relative to all other regions of the genome (i.e., normalized by genomic region size); and 2) how highly connected within protein networks are genes in the 22q11.2 locus to genes in disease-associated modules, relative to random gene-sets (i.e., normalized by number of seed genes)?

Relative to all regions of the genome (i.e., all Genomic-Region_BD-PPI_ networks), the 22q_BD-PPI_ network was above the 90^th^ percentile for 10 of 18 modules, including 6 of 7 SCZ- or ASD-associated modules (M1, M4, M6, M7, M13, M15; Fig. 3A, Table S4). Indeed, the percentile ranks of the 22q_BD-PPI_ network were significantly higher for SCZ- and/or ASD-associated modules than for non-SCZ- or ASD-associated modules, p=0.03, highlighting that relative to the rest of the genome, the 22q_BD-PPI_ network loads disproportionately onto SCZ- and ASD-associated modules.

**Figure 3.**
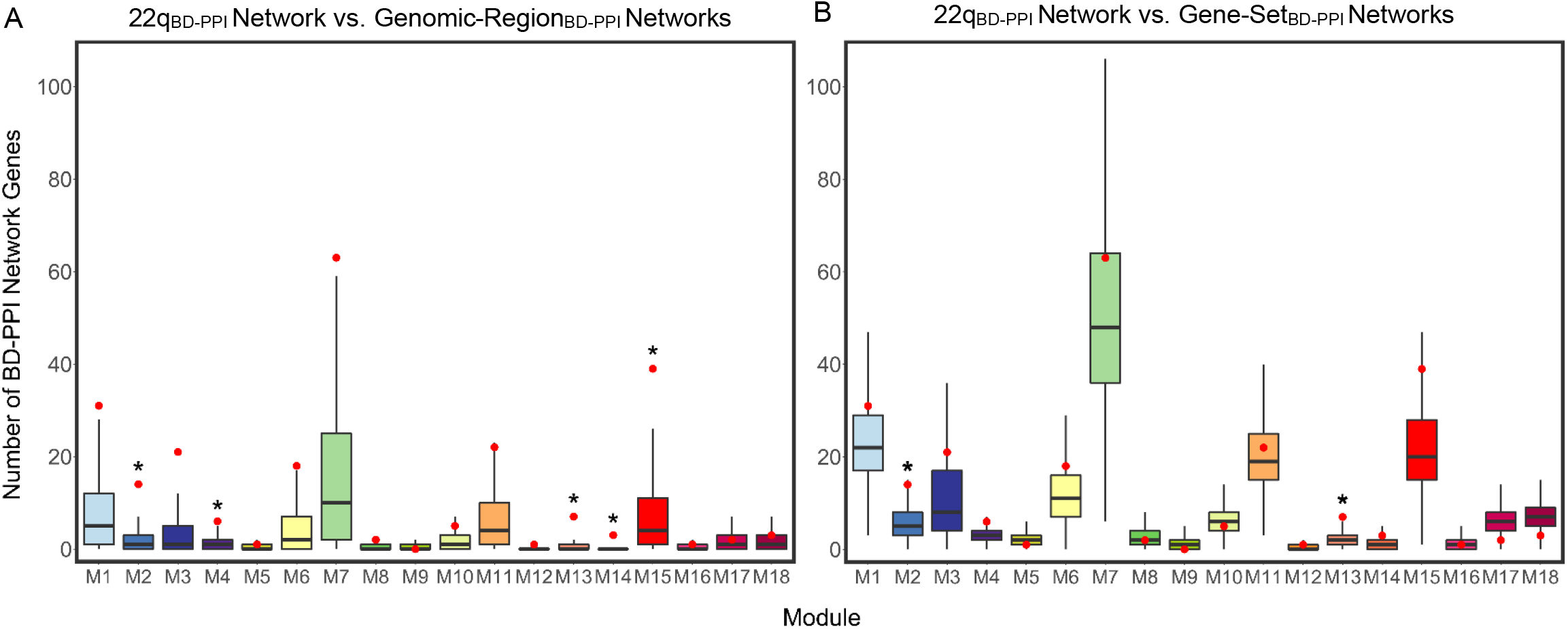
Characterizing the 22q_BD-PPI_ Network Relative to Genomic-Region and Gene-Set BD-PPI Networks. Distribution of number of BD-PPI network genes per module across all: (A) ~2.57 Mb genomic regions (i.e., 12,149 Genomic-Region_BD-PPI_ networks) and (B) random 38 seed gene gene-sets (i.e., 10,000 Gene-Set_BD-PPI_ networks) derived to characterize the 22q11.2 locus relative to all other regions of the genome, and the connectivity of 22q11.2 genes within brain development protein networks, respectively. Boxplots mark the median genes per module, with lower and upper hinges corresponding to the 25^th^ and 75^th^ quartiles, respectively, and the whiskers extending to 1.5 x interquartile range. The number of 22q_BD-PPI_ network genes in each module are marked in red. *22q_BD-PPI_ network above the 95th percentile.

Relative to the connectivity of random sets of 38 seed genes (i.e., the Gene-Set_BD-PPI_ networks), the 22q_BD-PPI_ network was above the 75^th^ percentile for 5 SCZ- and/or ASD-associated modules, including remaining in the ~96^th^ and ~93^rd^ percentiles of the SCZ- and ASD-associated M13 and M15 neuronal modules, respectively (Figure 3B; Table S4). The percentile ranks of the 22q_BD-PPI_ network relative to the Gene-Set_BD-PPI_ networks were similarly higher for SCZ- and/or ASD-associated modules than for non-SCZ- or ASD-associated modules, *p*=0.03.

Together, this suggests that 22q11.2 CNVs confer risk for SCZ and ASD by not only spanning a genomic region that is rich in protein-coding genes, but by spanning genes that form protein networks in the developing brain that are highly - and disproportionately - connected to broader networks associated with SCZ and ASD pathogenesis.

### *SEPT5, PI4KA*, and *SNAP29* are Candidate Drivers of Synaptic Pathology, and *DGCR8* and *HIRA* are Candidate Drivers of Dysregulated Gene Expression

To identify specific genes driving the 22q_BD-PPI_ network loading onto SCZ- and ASD-associated modules, we next examined the size of each individual 22q11.2 gene’s BD-PPI network for each module, relative to all BrainSpan-indexed genes. *SEPT5* and *PI4KA* were above the 99^th^ percentile for BD-PPI network genes in 3 neuronal modules associated with SCZ and ASD (M4, M13, M15; Fig. 4); *PI4KA* was additionally above the 95th percentile for the ASD-associated module, M12. *SNAP29* was also above the 95^th^ percentile for network genes in M13 and M15, as well as M7. Indeed, the individual *SEPT5, PI4KA*, and *SNAP29* networks were each enriched for cell parts and/or biological processes related to synaptic function (Fig. 5A-F). The *HIRA*_BD-PPI_ network was additionally above the 95^th^ percentile for genes in M7 and was enriched for genes involved in chromatin organization and gene expression (Fig. 5G,H), consistent with HIRA’s role as a histone chaperone protein. Additionally, the *DGCR8*_BD-PPI_ network was near the 99^th^ percentile of BrainSpan genes for M1 and was enriched for RNA binding and processing, consistent with *DGCR8’s* known role in microRNA (miRNA) biogenesis (Fig. 5I,J). Finally, the *MRPL40* and *ESS2* BD-PPI networks were above the 95^th^ percentile for M6; the *ESS2* network was not enriched for any GO terms, while *MRPL40*’s network was enriched for ribosomal components and mitochondrial elongation (Fig. 5K-M).

**Figure 4.**
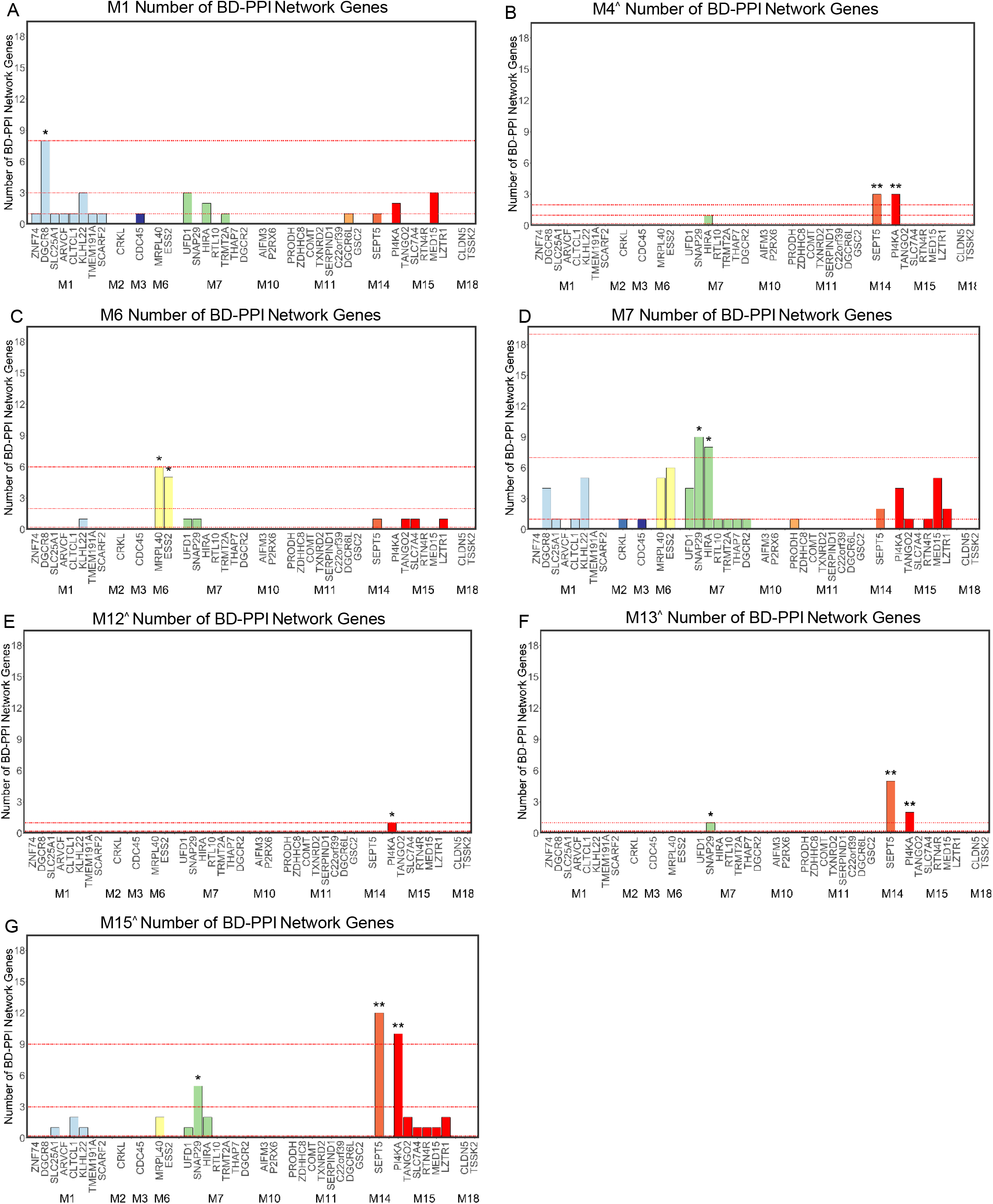
Prioritizing 22q11.2 Genes Based on BD-PPI Network Loading on BrainSpan Modules Associated with SCZ and ASD. Number of BD-PPI network genes per individual 22q11.2 gene in the: (A) M1, (B) M4, (C) M6, (D) M7, (E) M12, (F) M13, and (G) M15 SCZ and/or ASD-associated modules. 22q11.2 genes are shown grouped by module membership. The top, second, and third red dotted lines in each plot mark the number of BD-PPI network genes nearest to the 99^th^, 95^th^, and 75^th^ percentiles for each module. Genes above the 95^th^ or 99^th^ percentile for each module are denoted by * or **, respectively. ^Module enriched for neuronal cell-type markers.

**Figure 5.**
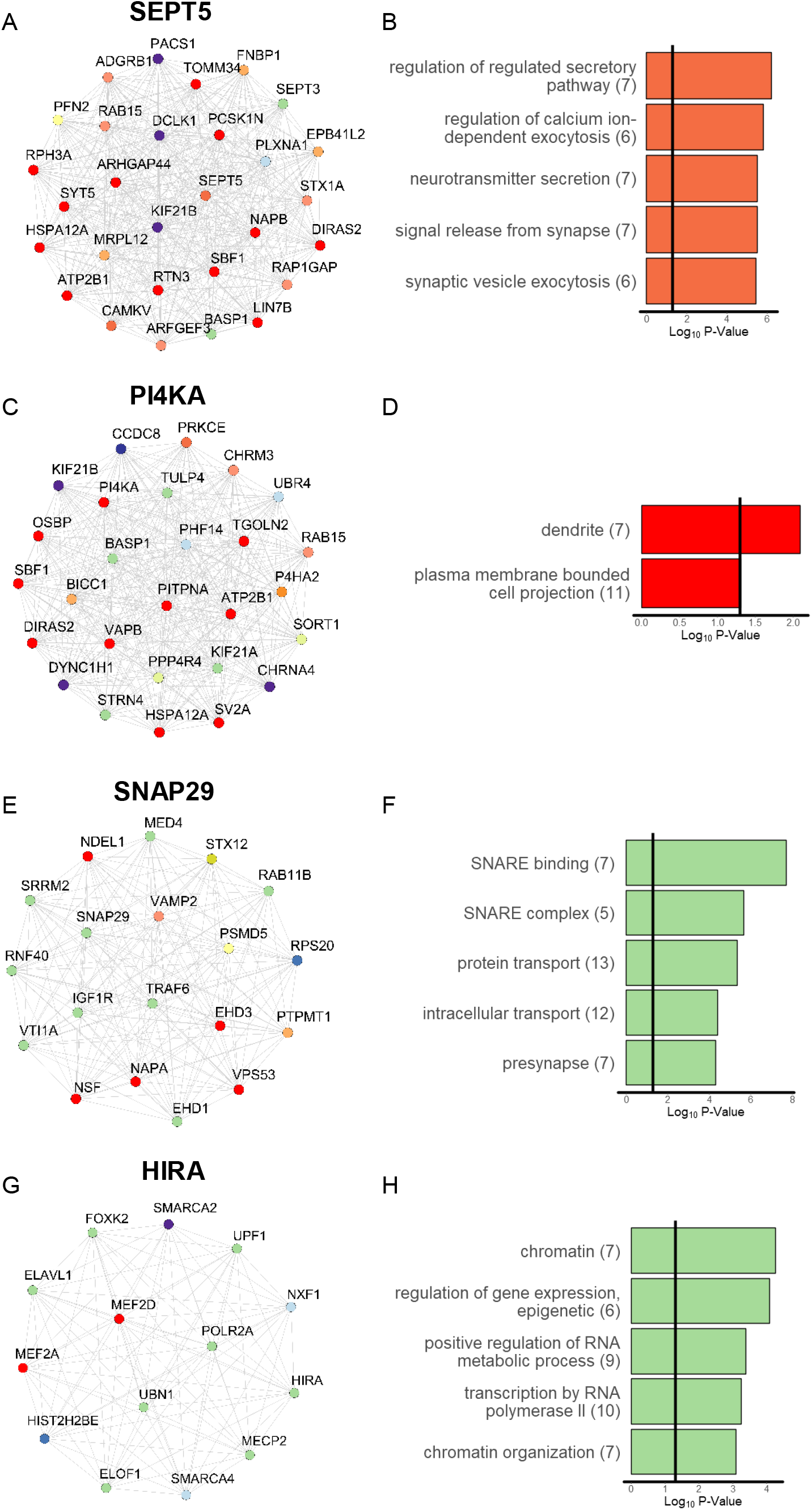
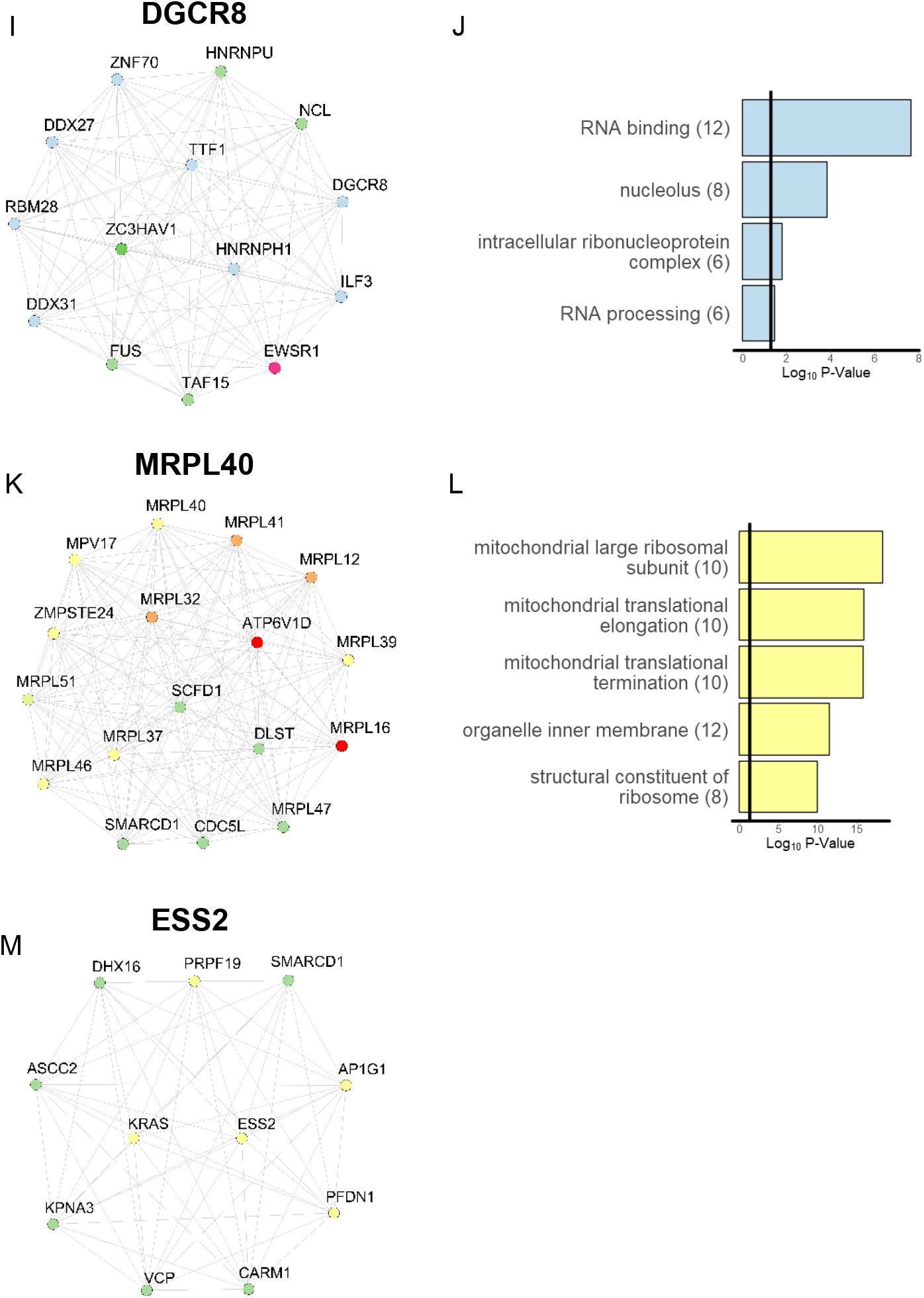
Characterization of BD-PPI Networks for 22q11.2 Genes Prioritized to Drive Risk for SCZ and/or ASD. Left: BD-PPI network genes and Right: significantly enriched GO terms (up to top 5 terms, where applicable) for the BD-PPI networks of: (A,B) SEPT5, (C,D) PI4KA, (E,F) SNAP29, (G,H) HIRA, (I,J) DGCR8, (K,L) MRPL40, and (M) ESS2. Gene nodes in each network are colored by primary BrainSpan module assignment.

### The 22q_BD-PPI_ Network Loads Highly onto SCZ and ASD Pathophysiologic Signatures

Post-mortem brain tissue from SCZ and ASD patients is usually ascertained from adults, limiting the detectability of differential gene expression (DGE) profiles that may exist during early brain development. Nevertheless, DGE profiles provide important pathophysiological signatures of disease. All modules enriched for genetic risk for SCZ were significantly enriched for down-regulated genes in SCZ cortex (M7, M13, M15; Fig. 6A). Similarly, 6 of 7 modules enriched for genetic risk for ASD were nominally or significantly enriched for down-regulated genes in ASD cortex (M4, M6, M7, M12, M13, M15). M1 was consistently enriched for ASD risk variants, but not for down-regulated genes in ASD cortex. However, M1 was highly enriched for genes expressed specifically during fetal development, suggesting that this likely reflects a floor effect in postnatal expression of these genes. Down-regulated genes in ASD and SCZ cortex therefore appear to at least partially reflect direct effects of genetic risk, and we focused on 22q_BD-PPI_ network overlap with down-regulated genes for subsequent analyses.

**Figure 6.**
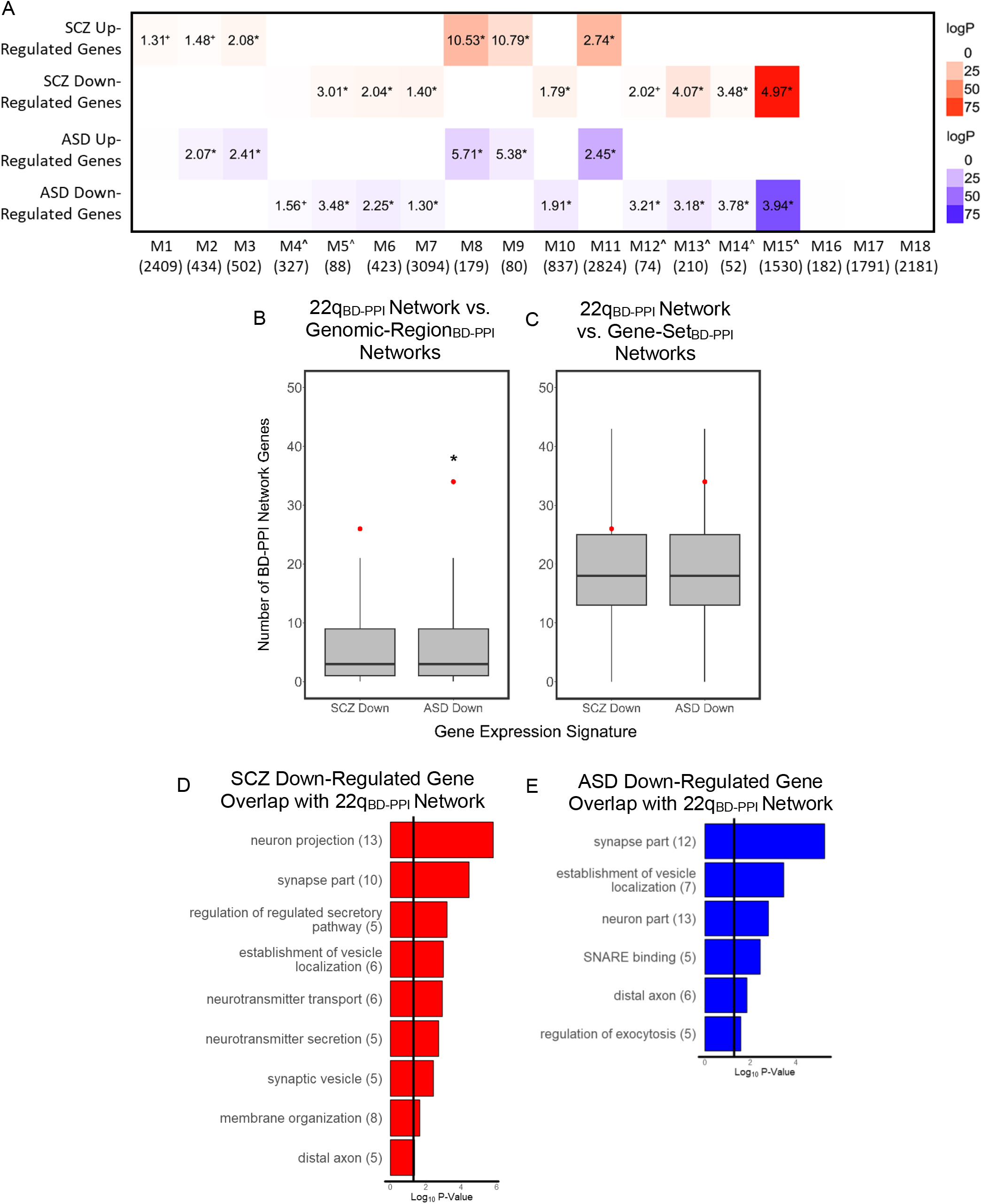
BrainSpan Module Enrichment and 22q11.2_BD-PPI_ Network Overlap for SCZ and ASD Pathophysiological Signatures. (A) BrainSpan module enrichment for significantly up- and down-regulated genes in SCZ (red) and ASD (blue) post-mortem cortex. Log p values are indicated by color for all enrichment odds ratios >1.0. Enrichment values for p<0.05 are annotated (+p<0.05, and *FDR corrected p<0.05 for number of modules), with numerical values denoting odds ratios for enriched modules. ^Module enriched for neuronal cell-type markers. Distribution of number of genes overlapping down-regulated genes in SCZ and ASD cortex for all: A) genome-wide ~2.57 Mb genomic regions (i.e., Genomic-Region_BD-PPI_ networks) and (B) random 38 seed gene gene-sets (i.e., Gene-Set_BD-PPI_ networks) derived to characterize the 22q11.2 region relative to all other regions of the genome, and the connectivity of 22q11.2 genes relative to all BrainSpan genes, respectively. Boxplots mark the median BD-PPI network genes per gene expression signature, with lower and upper hinges corresponding to the 25^th^ and 75^th^ quartiles, respectively, and the whiskers extending to 1.5 x interquartile range. The number of 22q_BD-PPI_ network genes for each gene signature are marked in red. *22q_BD-PPI_ network above the 95th percentile. Significantly enriched gene ontology (GO) terms for 22q_BD-PPI_ network genes overlapping down-regulated genes in (D) SCZ and (E) ASD cortex.

Given the size of the 22q_BD-PPI_ network and each DGE gene-set, the 22q_BD-PPI_ network was significantly enriched for down-regulated genes in both SCZ (~11% of the 22q_BD-PPI_ network; *p*=0.006) and ASD (~14% of the 22q_BD-PPI_ network; *p*=2.0×10^-4^). Additionally, compared to all regions of the genome (i.e., Genomic-Region_BD-PPI_ networks), the 22q_BD-PPI_ network was in the ~93^rd^ and ~96^th^ percentiles for overlap with down-regulated genes in SCZ and ASD, respectively (Fig. 6B; Table S4). Compared to the connectivity of random sets of 38 seed genes (i.e., Gene-Set_BD-PPI_ networks), the 22q_BD-PPI_ network was in the ~77^th^ and ~92^nd^ percentiles for down-regulated SCZ and ASD genes, respectively (Fig. 6C; Table S4). Notably, 22q_BD-PPI_ network genes overlapping down-regulated genes in SCZ and ASD were both enriched for neuronal and synaptic gene-sets (Fig. 6D,E), indicating that the 22q_BD-PPI_ network overlaps key pathophysiological signatures in SCZ and ASD via neuronal and synaptic features.

### *SEPT5* and *PI4KA* are Candidate Drivers of Synaptic Pathology Shared in SCZ and ASD

Among the 22q11.2 genes, *SEPT5* was above the 99^th^ percentile and *PI4KA* was near the 99^th^ percentile of BrainSpan genes for BD-PPI network overlap with down-regulated genes in SCZ and ASD (Fig. 7). Indeed, *SEPT5* accounted for ~50% and ~26% of the overall 22q_BD-PPI_ network overlapping down-regulated genes in SCZ and ASD cortex, respectively, and *PI4KA* accounted for ~37% and ~18% of the 22q_BD-PPI_ network overlap with down-regulated genes in SCZ and ASD, respectively. *SNAP29* and *KLHL22* were also above the 95^th^ percentile for BD-PPI network overlap with down-regulated genes in ASD, and accounted for ~15% and ~12% of 22q_BD-PPI_ network overlap with down-regulated genes in ASD, respectively. Thus, *SEPT5* and *PI4KA* were further highlighted as potential key drivers of synaptic pathology in 22q11.2 CNVs relevant to SCZ and ASD risk.

**Figure 7.**
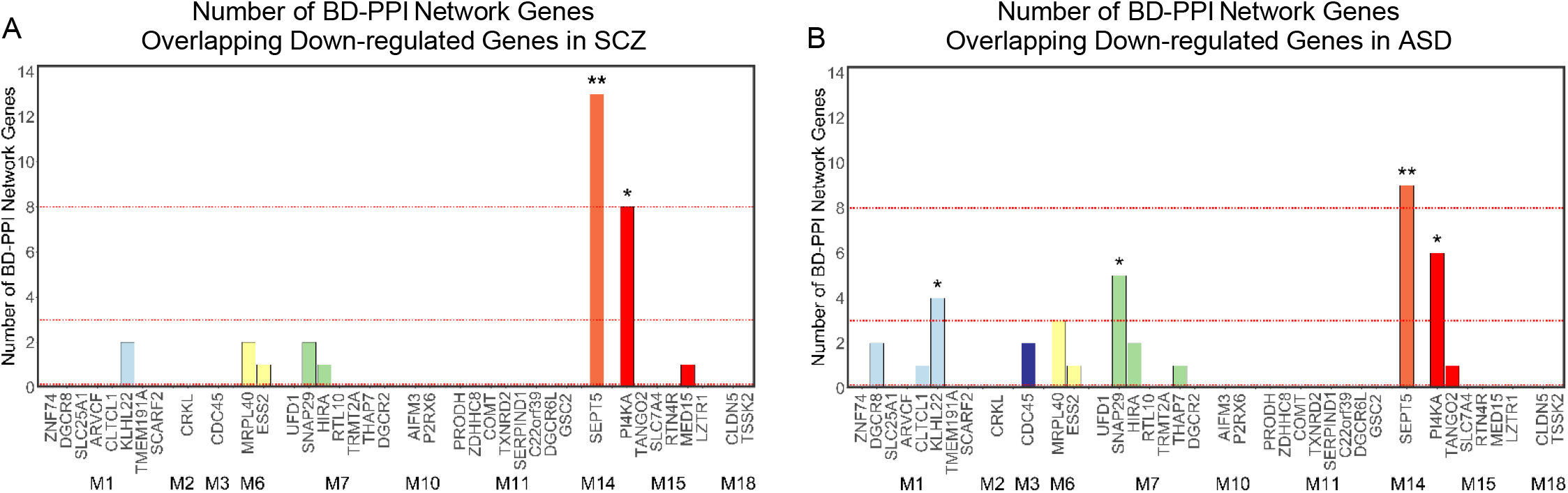
Prioritizing 22q11.2 Genes Based on BD-PPI Network Overlap with Key SCZ and ASD Pathophysiological Signatures. Number of BD-PPI network genes per individual 22q11.2 gene overlapping down-regulated genes in (A) SCZ and (B) ASD cortex. 22q11.2 genes are shown grouped by module membership. The top, second, and third red dotted lines on each plot mark the number of genes nearest to the 99^th^, 95^th^, and 75^th^ percentiles for all BrainSpan genes’ BD-PPI networks, when above 0. Genes above the 95^th^ or 99^th^ percentile for each gene expression signature are denoted by * or **, respectively.

### Supporting Analyses for the Current Framework for Understanding 22q11.2 CNV-Mediated Risk for SCZ and ASD

Exploratory analyses of broader CNVs and genes associated with SCZ and ASD confirmed that high BD-PPI network loading onto SCZ- and ASD-associated BrainSpan modules and down-regulated genes in SCZ or ASD cortex is a common characteristic of disease-associated CNVs and genes (Figs. S3 and S4, respectively; see Supplementary Information). Similarly, using lower co-expression thresholds (i.e., more inclusive thresholds) to define the BD-PPI networks of each gene highlighted similar 22q11.2 genes within the 22q11.2 locus as likely drivers of SCZ and ASD risk (Fig. S5). Furthermore, a machine learning approach to classify SCZ and ASD associated genes based on BD-PPI network percentile ranks across BrainSpan modules and DGE signatures prioritized similar 22q11.2 genes as likely disease-associated (Tables S1, S7). Finally, study bias could not account for our prioritization of 22q11.2 genes likely to drive risk for SCZ and ASD (Fig. S6). See Supplementary Information for details.

## Discussion

Recent genetic discoveries in SCZ and ASD have brought renewed focus to understanding the mechanisms through which genetic variation confers risk for these disorders at both the polygenic and individual variant level. Here, using a developmental functional genomics approach, we identified overlapping and distinct neurodevelopmental processes associated with risk for SCZ and ASD, and developed a framework leveraging this disease-informed lens to elucidate mechanisms underlying disease risk in highly penetrant multi-gene CNVs. Our analyses confirmed the capacity of 22q11.2 CNVs to disrupt disease-relevant neurodevelopmental networks and highlighted both well-studied (e.g., *DGCR8* and *HIRA*) and relatively understudied genes (e.g., *SEPT5, PI4KA*, and *SNAP29*) as candidate 22q11.2 genes likely to drive risk for SCZ and/or ASD.

Prior studies using transcriptomic data from the developing brain to understand how risk variants contribute to disease largely focused on rare variants in ASD. These studies suggested that ASD risk variants preferentially disrupt neuronal development and transcriptional and chromatin regulation during fetal development(15,16,34). One early study of DNMs in SCZ also implicated disruptions in neurogenesis and transcriptional regulation(25); however, this was a relatively small study. Here, by creating co-expression networks from brain samples spanning the full period of development and testing an expanded and updated set of risk variants for SCZ and ASD simultaneously, we confirmed prior enrichment patterns in ASD and identified co-expression modules with distinct enrichment patterns for ASD versus SCZ. We found robust enrichment in SCZ for a neuronal module that increased in expression across development and was involved broadly in postnatal synaptic transmission and refinement of neurotransmission; a neuronal module involved in regulating neuronal excitability that stabilized in expression around birth; and a module involved in regulating transcription and splicing across development. ASD risk variants showed overlapping enrichment for these modules, as well as robust enrichment for distinct modules involved in neuronal differentiation, and in regulating chromatin organization and gene transcription during fetal development. While our findings thus confirm prior findings implicating neuronal and synaptic genes in both SCZ and ASD, risk variants for ASD appear to partially diverge from those for SCZ by loading highly onto gene networks that orchestrate neuronal differentiation and synapse formation during fetal development. It is tempting to suggest that the higher loading of ASD risk variants onto early neuronal differentiation networks may be a key mechanism underlying the earlier onset of ASD relative to SCZ; however, experimental validation is necessary to confirm this.

By integrating BrainSpan co-expression and PPI data, we identified the protein network likely to be strongly affected by 22q11.2 CNVs during brain development. One prior study focused on 16p11.2 CNVs integrated PPI and BrainSpan co-expression data and found that ‘neuropsychiatric’ CNV networks were enriched for varying developmental periods, with the 22q11.2 network enriched during fetal and postnatal development(40). Here, using a disease-informed framework, we focused on understanding how 22q11.2 CNVs confer risk for SCZ and ASD, more specifically. We found that 22q11.2 genes formed a large BD-PPI network that: 1) loaded highly onto multiple SCZ- and ASD-associated modules, relative to all other regions of the genome; 2) was more highly connected to genes in SCZ- and/or ASD-associated modules than to genes in non-disease associated modules; and 3) overlapped highly with genes that are down-regulated in SCZ and ASD cortex, with overlapping genes enriched for neuronal and synaptic genes. This suggests that 22q11.2 CNVs may confer risk for SCZ and ASD, at least partially, by perturbing synaptic networks that modulate neuronal excitability and signaling. Consistent with this, mouse models of 22q11DS show altered neuronal excitability, dendritic spine stability, and synaptic activity(52,53).

A primary advantage of our approach was our ability to systematically survey 22q11.2 genes to prioritize genes likely to contribute to disease risk. Our analyses consistently highlighted *SEPT5* and *PI4KA* as candidate drivers of synaptic pathology relevant to SCZ and ASD, and *DGCR8* as a candidate driver of dysregulated gene expression that may be most relevant to ASD. *SNAP29* and *HIRA* were also prioritized as contributing to SCZ and ASD. Of note, both *SEPT5* and *SNAP29* have been found to negatively modulate neurotransmitter release. *SEPT5* encodes the filamentous protein septin 5, which is thought to serve as a molecular “brake” for neurotransmitter release by associating with synaptic vesicles to create a barrier that inhibits vesicle fusion with the membrane(55,56). Similarly, *SNAP29* encodes a SNARE protein that competes with α-SNAP for binding to SNARE complexes after membrane fusion, thus reducing SNARE protein recycling and synaptic vesicle availability(57). Conversely, altered *PI4KA* expression may contribute to SCZ and ASD risk by perturbing the proteomic composition of the plasma membrane (PM). *PI4KA* encodes the lipid kinase PI4KIIIα, which is involved in synthesizing the phosphoinositide, phosphatidylinositol 4,5-biphosphate (PI(4,5)P2), that recruits and regulates PM proteins and thereby governs diverse cellular processes including vesicular trafficking, ion transport, and actin cytoskeletal dynamics(58). *PI4KA* depletion was found to disrupt localization of PI(4,5)P2-associated proteins, reduce cellular responses to external stimuli(59, 60), and impair synapse maintenance(61). Our prioritization of *PI4KA* in contributing to SCZ and ASD risk is consistent with prior associations of *PI4KA* polymorphisms in SCZ(62–64) and with growing evidence that *PI4KA* may play an important role in synaptic function.

*DGCR8* is a core component of the microprocessor complex involved in processing miRNAs, which regulate gene expression at the protein level by binding target mRNAs to silence their translation(65). *HIRA* is a histone chaperone that influences gene expression by facilitating the deposition of the non-canonical histone variant, H3.3, into chromatin(66,67). Given these gene regulatory functions, *DGCR8* and *HIRA* have both been previously hypothesized to play an important role in 22q11.2 CNV phenotypes(67,68). Here, using our disease-informed functional genomic lens, we provide independent, unbiased evidence for the contribution of *DGCR8* and *HIRA* haploinsufficiency to risk for neurodevelopmental disorders.

Some caveats to the current study should be noted. First, in our analyses testing SCZ and ASD risk variants for module enrichment, some studies shared overlapping samples. As the genetic architectures of SCZ and ASD include many variants with small effects, collaborative analyses across studies are increasingly common to facilitate power. We utilized independent datasets where possible, and focused on modules showing consistent disease association.

Additionally, generating BD-PPI networks identifies protein networks putatively affected by 22q11.2 CNVs, but cannot tell us the functional output of increasing or reducing the expression of each gene. We therefore cannot determine how deletions vs. duplications of these genes affect broader cascades within these networks. Similarly, while using each gene’s BD-PPI network ranking is useful for prioritizing the most highly connected 22q11.2 genes within disease-associated modules, for genes whose mechanism of action involves regulating the expression of other genes, this cannot quantify the broader consequences of mutations in these genes. As high-throughput functional assays become increasingly feasible in the future, quantifying the transcriptomic and proteomic consequences of altered expression of each 22q11.2 gene will likely yield further insights into 22q11.2 CNV-mediated disease risk. Nevertheless, identifying convergent neurodevelopmental networks affected by polygenic risk for SCZ and ASD, and systematically prioritizing 22q11.2 genes that load highly onto these networks offers a powerful step towards understanding the specific mechanisms through which 22q11.2 CNVs confer risk for SCZ and ASD.

## Acknowledgements

The authors thank Gerhard Helleman for statistical advice and Elizabeth Ruzzo for feedback and advice on analyses. This work was supported by grants from the National Institutes of Mental Health (R01MH085953 to CEB, R01MH100900 to CEB, T32MH096682 to JKF, K08MH11877 to JKF).

## Conflicts of Interest

The authors have no conflicts of interest to disclose.

## Supplementary Methods

### Defining Modules of Developmentally Co-Expressed Genes

To identify modules of developmentally co-expressed genes in the human brain, co-expression networks were created from BrainSpan transcriptome microarray data(1), obtained from the Gene Expression Omnibus (https://www.ncbi.nlm.nih.gov/geo/query/acc.cgi?acc=GSE25219). BrainSpan is the largest, publicly-available developmental brain transcriptome dataset derived from 1340 post-mortem samples collected from 57 clinically unremarkable human donors, with no known psychiatric or neurologic conditions, aged 5.7 post-conception weeks (PCW) to late adulthood (82 years). Samples were collected across 16 brain regions, including 11 neocortical regions (see 1 for details).

Co-expression networks were created by conducting weighted gene co-expression network analysis (WGCNA; 2) using samples spanning the full period of brain development (i.e. 5.7 PCW – 30 years of age; 1061 samples). For genes indexed by multiple rows in GSE25219, the maximum mean expression value per gene was selected to represent the expression of each gene using the CollapseRows function, yielding expression measurements for 17220 unique genes with HGNC symbols. The majority of genes indexed in BrainSpan are protein-coding genes. Correlation coefficients were computed between the expression levels for each gene using biweight mid-correlation to create an adjacency matrix; a soft threshold of power of 15 was found to achieve approximate scale free topology (R^2^ > 0.85). The adjacency matrix was transformed into a topological overlap matrix (TOM) which sums the connection strength of each gene with every other gene. Genes were clustered with average linked hierarchical clustering based on the TOM to identify groups of co-expressed genes (i.e. modules). Modules were defined using the hybrid dynamic tree-cutting method(3) with standard parameters (minimum module size = 30, deepSplit = 2; 4). Modules were summarized by their first principal component (module eigengene, ME) and modules with ME correlations >0.9 were merged together. The final modules are annotated by a module number and corresponding color for illustrative purposes.

Genes in each module were functionally annotated using gene ontology (GO) biological and molecular pathways and cellular components from g:Profiler(5) with “moderate” hierarchical filtering (best term per parent) and a minimum query/term overlap size of 5 genes. Only pathways with 10 to 2000 genes were included in the analyses(6). A custom background was set to all genes indexed by BrainSpan. Top biological and molecular pathways surpassing g:SCS (Set Counts and Sizes) p<0.05 genes are shown for each module in the main results, with extended GO results including cellular components shown in Table S2.

Modules were additionally tested for primary brain cell-type enrichment using markers defined by(7), and for enrichment for the following gene-sets of interest derived from the literature on ASD and SCZ (i.e. previously found to be enriched for genetic variants associated with ASD and/or SCZ): chromatin modifiers(8), FMRP targets(9), RBFOX targets(10), Arc complex(10), NMDAR complex(10), PSD-95 complex(10), postsynaptic density(8), and loss-of-function intolerant genes (i.e. pLI genes;11). Enrichment was tested with Fisher’s exact tests, as implemented with the GeneOverlap package in R(12), with the total number of BrainSpan genes set as the genomic background. Modules were additionally tested for enrichment for lists of genes expressed in specific human brain regions during specific developmental periods, as well as for more specific brain cell sub-types, using the Specific Expression Analysis tool(13; http://genetics.wustl.edu/jdlab/csea-tool-2/), which provides graphical outputs summarizing the enrichment of input genelists at varying specificity thresholds.

### Identifying Neurodevelopmental Networks Associated with SCZ and ASD

Summary statistics, gene-lists, and/or variant lists from studies of common, rare, de novo, and copy number variants in SCZ and ASD were compiled from genome-wide association studies (GWAS) and tested for BrainSpan module enrichment using methods appropriate to each data type.

When summary level statistics for common and rare variants were available, they were tested for module enrichment using the default competitive gene-set enrichment model in MAGMA 1.06(14). MAGMA uses a multiple regression framework that aggregates genetic marker data to the level of genes using mean SNP p-values, and after accounting for linkage disequilibrium, tests the association between each trait and genes in a given gene-set compared to genes outside the gene-set (i.e. here, genes within each BrainSpan module vs. BrainSpan genes not in the module of interest). NCBI 37.3 gene definitions were used to restrict analyses to SNPs located within the transcribed region of each gene. For SCZ common variant GWAS, summary statistics from two studies were tested for enrichment, the PGC-deCODE meta-analysis(15) and the CLOZUK-PGC meta-analysis(16). Although these meta-analyses share overlapping subjects, summary statistics from both studies were examined as the SCZ CLOZUK-PGC meta-analysis includes 5,220 SCZ cases with a specific phenotype (i.e. treatment-resistant SCZ) and 18,823 controls that are not in the SCZ PGC-deCODE study, and the PGC-deCODE meta-analysis includes 1,513 SCZ cases and 36,989 controls from the deCODE study not in the CLOZUK-PGC study.

For de novo mutations (DNMs), individual validated DNMs from the literature were compiled for SCZ and ASD. The gene affected by each DNM and predicted functional consequence on the canonical transcript were annotated using Ensembl Variant Effect Predictor (VEP) version 90 for GRCh37(17). Indels predicted to have a frameshift consequence, and single nucleotide variants predicted to have stop-gain (i.e. nonsense), splice acceptor or splice donor consequences were defined as loss-of-function (LoF). LoF and missense variants were classified more broadly as protein-altering(18). Likelihood of the observed LoF and protein-altering mutation rate across patient groups for genes within each module were tested using a Poisson test against the expected rate, given gene-specific local sequence context (defined based on probability tables for the SSC sibling cohort from (19)) and implemented using the DenovolyzeR package in R(18). As negative control comparisons, the likelihood of the observed synonymous mutation rate in SCZ and ASD patients, and of synonymous, LoF, and protein-altering DNMs in control subjects derived from (20; 2196 control subjects) were also assessed per module, against the expected rate using DenovolyzeR.

For CNVs, HGNC gene symbols for SCZ- and ASD-associated loci were retrieved from Ensembl using the BioMart package in R; module enrichment for genes in SCZ- and ASD-associated CNVs was tested using logistic regression, controlling for gene length. Boundaries for SCZ-associated CNVs were defined based on (21; significant risk loci, FDR <0.05). (21) reported CNV borders in hg18; the UCSC LiftOver tool was used to convert them to hg19. ASD-associated CNVs were defined based on (22; significant risk loci in the combined SSC and AGP samples, FDR<0.05).

The CNV boundaries for six risk loci associated with SCZ and ASD shared more than 50% basepair (bp) overlap (see Table S3). For these 6 loci, the same CNV was considered to confer risk for both disorders and CNV boundaries were defined as the union between those provided by each study. For one additional locus (3q29), the coordinates associated with ASD in the combined SSC and AGP analysis overlapped ~28% with the SCZ defined locus; however, in the original analysis of the SSC sample alone, a larger 3q29 locus that overlapped 99% with the SCZ locus was significantly associated with ASD(22). We therefore considered the larger 3q29 locus to be a risk locus for both ASD and SCZ, and the union coordinates were used for both disorders for 3q29. Six additional loci were uniquely associated with SCZ and 5 additional loci were uniquely associated with ASD, 3 of which were nested within other ASD-associated loci.

Module enrichment for lists of genes containing more deleterious ultra-rare variants in SCZ patients than controls(10; p<0.01); manually curated high-confidence genes implicated in genetic studies of ASD according to the SFARI gene database (i.e. SFARI “Gene Score” of 1 or 2; data downloaded November 28, 2017; https://www.sfari.org/resource/sfari-gene/); and genes associated with ASD across rare inherited, de novo, and copy number variants (FDR≤0.1; 22) were formulated as a hypergeometric distribution and tested for significant overlap using Fisher’s exact tests, as implemented using the GeneOverlap package.

### Characterizing the 22q11.2 Locus and its Brain Development PPI Network

To first examine the protein-coding gene content of the 22q11.2 locus overall, all genes annotated as protein-coding in Ensembl were downloaded from BioMart in R. The number of protein-coding genes in the 22q11.2 locus was compared to a distribution generated by simulating CNVs of the same size (~2.57 mb) in 250 kb sliding steps across the entire genome, using the GenomicRanges package in R(23), yielding a set of 12,149 genome-wide simulated CNVs for comparison. A protein-coding gene was counted as within a simulated CNV region if any bp between the start and end site of the gene was located within the boundaries of the genomic region. The percentile rank of the 22q11.2 locus was calculated relative to this background distribution of simulated CNVs. To additionally examine whether any BrainSpan module was over-represented among the 22q11.2 genes indexed by BrainSpan, or whether SCZ- and ASD-associated modules were over-represented overall, a logistic regression controlling for gene length was conducted for each BrainSpan module, and for all disease-associated modules together, respectively.

To further elucidate why 22q11.2 CNVs confer elevated risk for SCZ and ASD, we sought to characterize the brain developmental protein network of the 22q11.2 locus relative to all other regions of the genome, and the connectivity of 22q11.2 genes within protein networks relative to all other genes, more specifically. We therefore compared the brain development protein-protein interaction network (BD-PPI network) of the 22q11.2 locus (22q_BD-PPI_ network) relative to those derived for: 1) all other equal-sized simulated CNV regions of the genome (i.e. the 12,149 ~2.57 Mb regions described above); and 2) 10,000 random gene-sets, equal in number of seed BrainSpan genes to the 22q11.2 locus (i.e., 10,000 x 38 BrainSpan gene gene-sets). To derive BD-PPI networks, a catalogue of physically interacting human proteins was first compiled by combining the BioGRID 3.4(24) and high-confidence InWeb3 PPI databases(25); after filtering these databases for PPI involving genes indexed in BrainSpan, this yielded a comprehensive catalogue of human PPI between BrainSpan genes containing 364,743 unique physical binary pairs. For each stage of brain development (stages 1-13 as defined in the original publication; 1), a catalogue of all highly co-expressed genes pairs was then generated from BrainSpan data by calculating the correlation between the expression level of each gene with every other gene, and retaining gene pairs co-expressed during each developmental stage at Spearman’s ρ>0.70(26). The comprehensive PPI catalogue was then thresholded by this catalogue of co-expressed gene pairs. Self-pairings and pairs that were redundant across multiple stages of development were removed, and each remaining gene-pair was flipped to capture the PPI relationship regardless of which gene in the pair was considered the “seed” gene. This yielded a comprehensive list of 77,832 bi-directional PPI pairings (38,916 unique pairs) between BrainSpan genes that were highly co-expressed during at least one stage of brain development. The mean number of PPI partners per BrainSpan gene in each module in the original PPI catalogue and after co-expression thresholding are shown in Table S5.

The 22q11.2 genes and their co-expressed PPI partners were extracted to identify the 22q_BD-PPI_ network. This was similarly done for the genes in all 12,149 genome-wide simulated CNV regions and 10,000 randomly generated, equal-length gene-sets, generating a catalogue of Genomic-Region_BD-PPI_ networks across the genome, and 10,000 Gene-Set_BD-PPI_ networks, respectively. The 22q_BD-PPI_ network was first tested for within-network enrichment for any BrainSpan modules, given the number of genes in each module, using hypergeometric overlap tests. The 22q_BD-PPI_ network was then characterized by its percentile rank for number of genes in each BrainSpan module compared to its corresponding Genomic-Region_BD-PPI_ networks and Gene-Set_BD-PPI_ networks. We focused on percentile ranks for these analyses given that these empirical distributions consist of the entire genome indexed by BrainSpan (i.e. including the 22q11.2 region and other SCZ- and ASD-associated loci). Mann-Whitney U tests were used to examine whether the percentile ranks for the 22q_BD-PPI_ network, relative to its corresponding Genomic-Region_BD-PPI_ and Gene-Set_BD-PPI_ networks, were higher overall for SCZ- and ASD-associated modules than for non-SCZ- or ASD-associated modules.

### Characterizing the BD-PPI Networks of Genome-Wide SCZ- and ASD-Associated CNVs

As an exploratory analysis, we also examined whether SCZ- and ASD-associated CNVs as a class are characterized by spanning genomic regions that generate BD-PPI networks that load highly onto at least one SCZ- or ASD-associated module, respectively, and/or span genes that are highly connected in protein networks with genes in at least one SCZ- or ASD-associated module. Thus, for each SCZ and/or ASD-associated CNV (Table S3) that spanned at least one BrainSpan gene, we simulated corresponding CNVs of the same size (range: ~0.17-4.76 Mb) in 250 kb sliding steps across the genome, and 10,000 gene-lists consisting of the same number of seed BrainSpan genes (range: 1-26 genes), parallel to the process for the 22q11.2 locus. We identified the BD-PPI networks for each set of genomic ranges and gene-sets, and then calculated the percentile rank of each SCZ- or ASD-associated CNV BD-PPI network relative to its corresponding Genomic-Region_BD-PPI_ and Gene-Set_BD-PPI_ networks for number of genes in each BrainSpan module.

To additionally generate a distribution for a set of more specific “control” CNV regions for comparison, we downloaded all CNVs from dbVar that were found in humans and categorized as “likely benign” or “benign” (hereaft er, benign; https://www.ncbi.nlm.nih.gov/dbvar/). We filtered the data for those CNVs that were between 0.25 – 5.0 Mb. This size range was chosen to include benign CNVs that are likely to be large enough to span genes, while remaining relatively close to the range of CNV sizes associated with SCZ and ASD. We then used the disjoin function from the GenomicRanges package to find all non-overlapping genomic ranges represented in this set of benign CNVs. We retained all regions spanned by at least 2 benign CNVs to increase the likelihood that CNVs in those regions have truly benign consequences. To generate a distribution for the maximum percentile rankings of Benign CNV BD-PPI networks on SCZ- and ASD-associated modules, we first identified the 20 largest regions spanned by benign CNVs in dbVar (size range: ~3.40-0.55 Mb; Table S6). For each region, parallel to the process for SCZ- and ASD-associated CNVs, we identified the BD-PPI networks for simulated CNVs of the same size in 250 kb sliding steps across the genome and calculated the percentile ranking of the Benign CNV BD-PPI network on each BrainSpan module relative to their corresponding distribution of Genomic-Region_BD-PPI_ networks. As many of the benign CNV regions did not contain any BrainSpan genes, to retain a large enough sample of benign CNVs to characterize the connectivity of BrainSpan genes in protein networks when they do exist in benign CNVs, we then identified the 20 largest regions containing at least 1 BrainSpan gene (range: 1 to 19 genes; Table S6) and generated 10,000 gene-lists consisting of the same number of seed BrainSpan genes for each region. We similarly identified the BD-PPI network for each gene-set and calculated the percentile rank of the Benign CNV BD-PPI network on each BrainSpan module relative to their corresponding Gene-Set_BD-PPI_ networks.

Given our hypothesis that SCZ- and ASD-associated CNVs confer risk by disrupting at least one disease-associated neurodevelopmental process, we extracted the maximum percentile rank of each BD-PPI network across the SCZ-associated (M7, M13, M15) and non-SCZ-associated modules, and across the ASD-associated (M1, M4, M6, M7, M12, M13, M15) and non-ASD-associated modules. Mann-Whitney U tests were used to compare the maximum percentile ranks of SCZ-associated CNV BD-PPI networks for SCZ-associated and non-SCZ-associated modules versus the distribution generated by concatenating the maximum percentile ranks of their corresponding Genomic-Region_BD-PPI_ networks and Gene-Set_BD-PPI_ networks, and the distribution for Benign CNVs. Similarly, the maximum percentile ranks of ASD-associated CNV BD-PPI networks for ASD-associated and non-ASD-associated modules were compared to the distribution for their corresponding Genomic-Region_BD-PPI_ networks and Gene-Set_BD-PPI_ networks, and the Benign CNVs.

### Characterizing the BD-PPI Networks of Genome-Wide SCZ- and ASD-Associated Genes

To strengthen our conclusion that individual 22q11.2 genes with BD-PPI networks that load highly onto SCZ- or ASD-associated modules are likely to drive disease risk, we assessed whether genome-wide SCZ- and ASD-associated genes form BD-PPI networks with higher maximum percentile ranks for SCZ- and ASD-associated modules, respectively, compared to all other BrainSpan genes and to “control” genes found in the Benign CNVs focused on in the above analyses (86 genes from CNVs listed in Table S6).

Genes identified as significantly associated with SCZ and ASD were compiled from the datasets listed in Table 1. For SCZ (472 unique BrainSpan genes), the vast majority of genome-wide significant genes were identified from the MAGMA based analyses of common variants in SCZ (Bonferroni p<0.05; corrected for 20,000 genes). Genes in CNV loci that were significantly associated with SCZ and only spanned a single gene (i.e. NRXN1, DMRT1) were also included. For ASD (109 unique BrainSpan genes), all SFARI genes, 65 genes identified in (22), genes with genome-wide significant enrichment of LoF or protein-altering DNMs, single gene CNVs (i.e. NRXN1), and genome-wide significant genes identified from the MAGMA-based analysis of common variants in ASD were included. Genes utilized as SCZ- or ASD-associated are annotated in Table S1. We note that SHANK3 is the only gene in the 22q13.3 ASD-associated CNV locus; however, it was not indexed in the BrainSpan microarray data, and thus was not included in these analyses. Mann-Whitney U tests were used to compare the maximum percentile ranks of SCZ-associated gene BD-PPI networks for SCZ-associated and non-SCZ-associated modules compared to the BD-PPI networks for all genes in BrainSpan not significantly associated with SCZ, and for “control” genes found in the Benign CNVs. We also compared the BD-PPI networks of ASD-associated genes to non-ASD-associated genes and “control” genes for ASD-associated and non-ASD-associated modules. Of note, given that the “non-SCZ-associated” and “non-ASD-associated” gene groups in these analyses were defined broadly as any BrainSpan gene not significantly associated with SCZ or ASD, respectively, as described above, many of these genes may be found to be associated with SCZ and ASD in future studies, as psychiatric genomic studies continue to gain statistical power. The “non-SCZ-associated” and “non-ASD-associated” gene groups also included genes spanned by multi-gene CNV loci associated with SCZ and ASD, as it is unclear at this time which genes within these loci are pathogenic. These “non-SCZ-associated” and “non-ASD-associated” gene groups can therefore be considered conservative comparison groups that are complementary to the “control” group of genes found in Benign CNVs.

Parallel analyses were conducted for the BD-PPI network percentile ranks of SCZ-associated genes for down-regulated genes in SCZ post-mortem cortex, and ASD-associated genes for down-regulated genes in ASD post-mortem cortex.

### Machine Learning Prediction of SCZ- and ASD-Associated Genes

To determine if an alternative, machine learning approach would prioritize similar 22q11.2 genes as conferring risk for SCZ and ASD, we applied a linear kernel support vector machine (SVM), implemented with caret in R(27), to predict disease association using BD-PPI network ranks across BrainSpan modules and down- and up-regulated genes in SCZ or ASD, respectively (i.e. 20 features per disease model). To train our models for SCZ and ASD, we used our lists of genome-wide SCZ-(472 genes) and ASD-(109 genes) associated genes, respectively, as the “positive” class, and an extended list of “control” genes found in benign CNV regions as the “negative” class. One 22q11.2 gene was significantly associated with SCZ in MAGMA-based analyses of common variants (ZDDHC8;16); this gene was therefore included in the training data and was not examined during the prediction step. For the extended list of benign CNV “control” genes, we used genes spanned by any of the 0.25-5.0 Mb benign CNV regions downloaded from dbVar and found in at least 2 individuals. Of 233 unique genes identified from this process, one was significantly associated with ASD based on analyses of common variants in ASD (KANSLI) and 11 additional genes were spanned by CNV regions associated with ASD or SCZ. These genes were dropped, leaving 221 unique genes to include as “control” genes.

Nested *K*-fold cross validation (CV) with *k* = 10 was used to optimize the regularization parameter (C) for predictive accuracy and estimate the accuracy of the best performing model across each fold. Thus, the original training data for each disorder was divided into 10 balanced folds of 90/10 training and test splits for the outer CV loop. The training split of each fold was then input into an inner 10-fold CV loop to optimize *C* using the held-out samples of each inner fold; the optimal *C* was used to train the final model for each fold and applied to the test split of each fold. The Synthetic Minority Over-sampling Technique (SMOTE; 28) algorithm was applied to account for class imbalance in the training data. The overall accuracy of the model selection procedure was calculated by comparing the predictions of the test split of each fold to the original class designations and tested for significance using a one-sided binomial test relative to the “no information rate” model, based on the largest class percentage in the data(27). To generate stable SCZ and ASD class predictions for the 22q11.2 genes, as well as for all BrainSpan genes not in the training data, the final model derived from each fold of the training data was applied to all other BrainSpan genes; the probability of SCZ or ASD association for each gene was averaged across the final 10 models (i.e. based on each of the 10 folds) and is reported in Table S1.

### Examining the Potential Relationship Between Study Bias and Prioritized 22q11.2 Genes

Analyses incorporating PPI data can be biased towards well-studied genes, as the PPI networks of well-studied genes may be characterized more completely in the literature than that for less-studied genes. To examine whether study bias could account for our prioritization of genes within the 22q11.2 locus, we therefore obtained the number of PubMed papers citing each 22q11.2 gene indexed in BrainSpan as of October 7, 2018, and examined the correlation (Spearman’s ρ) between number of published papers per gene and the number of BD-PPI network partners across all SCZ- and ASD-associated modules; maximum percentile rank of BD-PPI network partners in any SCZ- or ASD-associated module; and number of BD-PPI network partners overlapping down-regulated genes in SCZ or ASD(29).

## Supplementary Results

### BrainSpan Module Enrichment for Literature Based Gene-Sets Are Consistent with GO Analyses

All modules significantly associated with SCZ or ASD were enriched for LoF intolerant (pLI) genes (Fig. S1C). Module enrichment for gene-sets implicated in prior genetic studies of SCZ or ASD were consistent with GO analyses, with M1 and M7 enriched for chromatin modifiers, M4, M13, and M15 enriched for postsynaptic density (PSD) genes, and M13 and M15 additionally enriched for the NMDAR, Arc, and PSD-95 complexes (Fig. S1C). M4, M6, M7, M12, M13, and M15 were also enriched for RBFOX targets, and M4, M12, M13, and M15 were enriched for FMRP targets.

### SCZ- and ASD-Associated CNV BD-PPI Networks Load Highly onto Disease-Associated Modules

To strengthen our conclusion that the high loading of the 22q_BD-PPI_ network onto SCZ- and ASD-associated networks is likely to reflect disease-relevant mechanisms of 22q11.2 CNVs, we asked whether SCZ- and ASD-associated CNVs, as a class, span genomic regions that generate BD-PPI networks that load highly onto at least one disease-associated module and/or contain genes that are highly connected within brain development protein networks to genes in disease-associated modules.

Normalized by genomic region size, the BD-PPI networks of SCZ-associated CNVs had significantly higher maximum rank percentiles for both SCZ- and non-SCZ-associated modules compared to their corresponding Genomic-Region_BD-PPI_ networks, and compared to Benign CNV BD-PPI networks (Fig. S3A,B). However, normalized by number of seed genes (i.e. Gene-Set_BD-PPI_ networks), SCZ CNV BD-PPI networks had significantly higher maximum percentile ranks than Benign CNV BD-PPI networks only for SCZ-associated modules (Fig. S3C,D). SCZ CNVs did not differ significantly from their corresponding Gene-Set_BD-PPI_ networks for SCZ- or non-SCZ associated modules. Thus, as a class, SCZ-associated CNV regions generate BD-PPI networks that load more densely onto both SCZ- and non-SCZ-associated modules compared to Benign CNV regions and the “average” region of the genome; however, their genes are more highly connected within brain protein networks to genes in SCZ-associated modules, specifically, compared to Benign CNVs.

Normalized by genomic region size, ASD CNV BD-PPI networks also showed significantly higher maximum percentile ranks for ASD- and non-ASD associated modules compared to Benign CNV BD-PPI networks; however, they had significantly higher rank percentiles than their corresponding Genomic-Region_BD-PPI_ networks only for ASD-associated modules (Fig. S3E,F). Normalized by number of seed genes, ASD CNV BD-PPI networks had significantly higher maximum rank percentiles than Benign CNV BD-PPI networks and their corresponding Gene-Set_BD-PPI_ networks specifically for ASD-associated modules (Fig. S3G,H). Thus, ASD-associated CNV BD-PPI networks load more highly onto ASD-associated modules than the “average” region of the genome, and also contain genes that are more highly connected within brain development protein networks to genes in ASD-associated modules than the “average” gene.

Together, this suggests that the high loading of the 22q_BD-PPI_ network onto multiple SCZ- and ASD-associated modules is likely to reflect disease-relevant mechanisms.

### SCZ- and ASD-Associated Gene BD-PPI Networks Load Highly onto Disease-Associated Modules

To verify the utility of leveraging the percentile rank loading of each gene’s BD-PPI network onto disease-associated modules and key pathophysiological signatures in order to pinpoint 22q11.2 genes relevant for ASD and SCZ risk, we compared the maximum percentile rank of ASD- and SCZ-associated genes to that for genes found in benign CNVs (i.e. “control” genes derived from Benign CNVs listed in Table S6; 86 genes) and for all BrainSpan genes not significantly associated with ASD (17111 genes) or SCZ (16748 genes), respectively.

The maximum BD-PPI network percentile rank across SCZ-associated modules was significantly higher for SCZ-associated genes than for “control” genes and non-SCZ-associated genes (Fig. S4A); SCZ-associated genes also had higher maximum percentile ranks than “control” genes for non-SCZ associated modules, but similar percentiles compared to non-SCZ-associated genes (Fig. S4B). SCZ-associated genes also had significantly higher BD-PPI network rank percentiles for overlap with down-regulated genes in SCZ post-mortem cortex compared to non-SCZ-associated genes and “control” genes (Fig. S4C).

Similarly, the maximum BD-PPI network rank percentiles for ASD-associated genes across ASD-associated modules was significantly higher than for “control” genes and non-ASD-associated genes (Fig. S4D). ASD-associated genes did not differ from “control” genes for non-ASD-associated modules, and had significantly lower percentile ranks than non-ASD-associated genes for non-ASD-associated modules (Fig. S4E). Finally, ASD-associated genes also showed significantly higher BD-PPI network rank percentiles for overlap with down-regulated genes in ASD cortex compared to non-associated genes and “control” genes (Fig. S4F).

Together, this suggests that identifying individual genes with BD-PPI networks that load highly onto modules of interest and key pathophysiological signatures is a useful strategy to identify individual genes likely to confer risk for these disorders.

### Robustness of Prioritized Disease-Relevant 22q11.2 Genes at Lower Expression Thresholds and Using a Machine Learning Approach

Using lower co-expression thresholds (i.e. more inclusive thresholds; Spearman’s ρ>0.50 and ρ>0.60) to define the BD-PPI networks of each gene highlighted similar genes as likely drivers of SCZ and ASD risk in 22q11.2 CNVs. Thus, the BD-PPI networks for SEPT5 and PI4KA were consistently above the 95th percentile across multiple SCZ- and/or ASD-associated neuronal modules, and for overlap with down-regulated genes in SCZ and ASD post-mortem cortex (Figs. S5). The percentile rank of SNAP29 for the M13 and M15 neuronal modules ranged from ~90-97%, and remained near the 95th percentile for the M7 gene regulatory module. DGCR8 was consistently above the 95th percentile for BD-PPI network overlap with M1. At lower BD-PPI network co-expression thresholds, the percentile rank of HIRA for M7 ranged from ~88-91%.

Additionally, while we focused on genes that formed BD-PPI networks above the 95th percentile for disease-associated modules, with the assumption that they may exert the most disruptive effects, we recognize that this a relatively arbitrary threshold and many genes with lower network loading on disease-associated modules may contribute risk. To determine if a machine-learning approach would prioritize similar 22q11.2 genes, we applied a support vector machine (SVM) to our lists of genome-wide SCZ- and ASD-associated genes and an extended list of benign CNV “control” genes, to predict disease association based on each genes’ BD-PPI network rankings across modules and gene expression signatures. Individual genes and variants contribute to SCZ and ASD risk with varying effect size; as our data inputs were binarized at the gene level, we could not account for this and our models could be interpreted as predicting disease-associated genes based on variants in these genes of similar effect size to the model inputs which came disproportionately from DNM studies for ASD (i.e., likely larger effects) versus common variants for SCZ (i.e., likely smaller effects). Nevertheless, our models significantly predicted ASD (Test Accuracy = 76.97%, *p* = 4.52 x10^-05^) and SCZ association class (Test Accuracy = 72.44%, *p* = .0075; Table S7). The ASD models identified 17 22q11.2 genes as predicted ASD-associated, including all genes highlighted in the above results for ASD, with the highest probabilities for *SEPT5* (97.32%), *HIRA* (97.04%), and *PI4KA* (96.97%). The SCZ models showed less discrimination between 22q11.2 genes, retaining 34 tested genes as predicted SCZ-associated; however, *PI4KA* (86.56%) and *SEPT5* (82.07%) showed the highest probabilities, followed by *TRMT2A* (80.83%), *HIRA* (80.12%), and *SNAP29* (79.06%). Table S1 displays the probabilities of SCZ and ASD association for all other BrainSpan genes.

### Study Bias Does Not Account for 22q11.2 Gene Prioritization

We found no significant relationships between number of published papers per gene and the number of BD-PPI network partners across all SCZ- and ASD-associated modules; maximum percentile rank of BD-PPI network partners in any SCZ- or ASD-associated module; or number of BD-PPI network partners overlapping down-regulated genes in SCZ or ASD (Fig. S6). Thus, study bias does not appear to account for our prioritization of 22q11.2 genes.

## Supplemental Tables and Figures

**Table S1. BrainSpan Gene Annotation.** BrainSpan genes are annotated by module assignment, localization to disease-related or benign CNV regions (where applicable), brain developmental protein-protein interaction (BD-PPI) network percentile rank for each module, and predicted probability of SCZ or ASD association based on linear SVM models.

**Table S2. Extended Gene Ontology (GO) Results for BrainSpan Modules.**

**Table S3.**
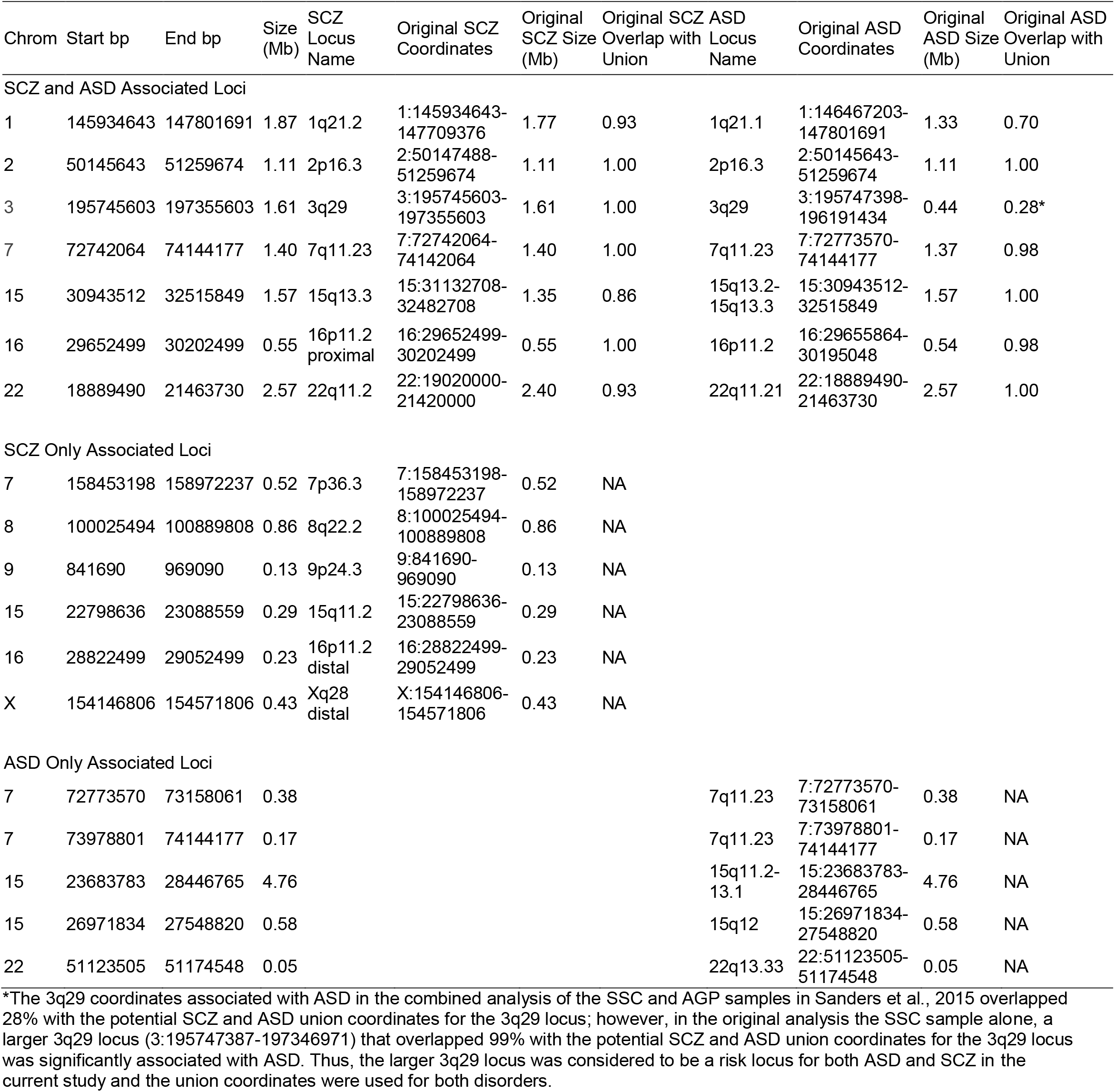
SCZ- and ASD-Associated CNV Loci. For loci associated with both SCZ and ASD (i.e., the individual SCZ or ASD associated loci overlapped and shared more than 50% of their respective bp with the potential union) the union coordinates between the individual SCZ and ASD associated loci were used for the current study.

**Table S4.** Percentile Rank Summary for 22q11.2 Genes and 22q11.2_BD-PPI_ Network for BrainSpan Modules and Gene Expression Signatures. Percentile rank and corresponding rank *p* value for number of 22q11.2 genes and 22q_BD-PPI_ network genes in each BrainSpan module and for overlap with down-regulated genes in SCZ and ASD post-mortem cortex compared to all simulated CNVs of the same size across the genome (Genomic-Region_BD-PPI_ networks), and to random lists generated from an equal number of BrainSpan seed genes (38 seed genes; Gene-Set_BD-PPI_ networks). Rank *p* values in the top 5% are denoted by *. ^Module enriched for neuronal cell-type markers.

|  | Number of 22q11.2 Genes | Percentile Rank for 22q11.2 Genes vs. Genomic Regions | Rank *P* for 22q11.2 Genes vs. Genomic Regions | Number of 22q<sub>BD-PPI</sub> Network Genes | Percentile Rank for 22q<sub>BD-PPI</sub> Network vs. Genomic-Region<sub>BD-PPI</sub> Networks | Rank *P* for 22q<sub>BD-PPI</sub> Network vs. Genomic-Region<sub>BD-PPI</sub> Networks | Percentile Rank for 22q<sub>BD-PPI</sub> Network vs. Gene-Set<sub>BD-PPI</sub> Networks | Rank *P* for 22q<sub>BD-PPI</sub> Network vs. Gene-Set<sub>BD-PPI</sub> Networks |
| --- | --- | --- | --- | --- | --- | --- | --- | --- |
| Module |  |  |  |  |  |  |  |  |
| M1 | 8 | 95.76 | 0.0424* | 31 | 94.29 | 0.0571 | 79.44 | 0.2056 |
| M2 | 1 | 73.2 | 0.268 | 14 | 98.52 | 0.0148* | 94.73 | 0.0527 |
| M3 | 1 | 70.06 | 0.2994 | 21 | 93.81 | 0.0619 | 79.21 | 0.2079 |
| M4^ | 0 | 0 | 1 | 6 | 96.17 | 0.0383* | 85.56 | 0.1444 |
| M5^ | 0 | 0 | 1 | 1 | 60.02 | 0.3998 | 17.66 | 0.8234 |
| M6 | 2 | 93.31 | 0.0669 | 18 | 94.1 | 0.059 | 80.6 | 0.194 |
| M7 | 7 | 91.07 | 0.0893 | 63 | 92.63 | 0.0737 | 72.38 | 0.2762 |
| M8 | 0 | 0 | 1 | 2 | 77.19 | 0.2281 | 36.76 | 0.6324 |
| M9 | 0 | 0 | 1 | 0 | 0 | 1 | 0 | 1 |
| M10 | 2 | 83.19 | 0.1681 | 5 | 83.08 | 0.1693 | 32.06 | 0.6794 |
| M11 | 8 | 93.91 | 0.0609 | 22 | 91.46 | 0.0854 | 60.74 | 0.3926 |
| M12^ | 0 | 0 | 1 | 1 | 79.86 | 0.2014 | 54.25 | 0.4576 |
| M13^ | 0 | 0 | 1 | 7 | 98.86 | 0.0114* | 95.99 | 0.0401* |
| M14^ | 1 | 95.41 | 0.0459* | 3 | 95.68 | 0.0432* | 88.36 | 0.1164 |
| M15^ | 6 | 96.27 | 0.0373* | 39 | 96.3 | 0.0370* | 92.56 | 0.0744 |
| M16 | 0 | 0 | 1 | 1 | 74.14 | 0.2586 | 32.48 | 0.6752 |
| M17 | 0 | 0 | 1 | 2 | 56.81 | 0.4319 | 2.29 | 0.9771 |
| M18 | 2 | 70.71 | 0.2929 | 3 | 72.79 | 0.2721 | 5.02 | 0.9498 |
| All Modules | 38 |  |  | 239 |  |  |  |  |
| Gene Expression Signature |  |  |  |  |  |  |  |  |
| SCZ Down | 0 | 0 | 1 | 26 | 93.14 | 0.0686 | 77.18 | 0.2282 |
| ASD Down | 3 | 88.04 | 0.1196 | 34 | 96.08 | 0.0392* | 92.12 | 0.07881 |

**Table S5.** Mean Connectivity of Genes in Each BrainSpan Module. Mean number of original protein-protein interaction (PPI) partners for genes in each BrainSpan module, based on the BioGrid and InWeb3 PPI databases, and mean number of brain developmental PPI (BD-PPI) partners, representing PPI partners that were highly co-expressed (Spearman’s ρ > 0.70) during at least one developmental stage in the human brain based on BrainSpan data. ^Module enriched for neuronal cell-type markers.

| Module | Number of Genes | Mean Number of Original PPI Partners (SD) | Mean Number of BD-PPI Partners (SD) |
| --- | --- | --- | --- |
| M1 | 2409 | 49.08 (82.34) | 4.03 (9.32) |
| M2 | 434 | 54.22 (71.97) | 6.04 (13.17) |
| M3 | 502 | 75.22 (109.28) | 13.47 (22.25) |
| M4^ | 327 | 32.43 (66.37) | 4.03 (12.02) |
| M5^ | 88 | 53.11 (75.9) | 11.66 (18.47) |
| M6 | 423 | 79.25 (107.18) | 14.53 (21.15) |
| M7 | 3094 | 65.49 (90.19) | 7.88 (14.5) |
| M8 | 179 | 61.7 (104.05) | 6.3 (10.91) |
| M9 | 80 | 55.06 (72.08) | 7.76 (13.4) |
| M10 | 837 | 33.2 (45.83) | 2.77 (7.09) |
| M11 | 2824 | 35.29 (69.18) | 2.65 (7.74) |
| M12^ | 74 | 25.31 (48.17) | 3.5 (8.57) |
| M13^ | 210 | 36.76 (49.26) | 4.28 (6.75) |
| M14^ | 52 | 46.71 (54.17) | 9.37 (14.24) |
| M15^ | 1530 | 47.03 (87.59) | 6.4 (14.23) |
| M16 | 182 | 32.94 (68.8) | 2.6 (5.98) |
| M17 | 1791 | 16.05 (39.23) | 0.74 (2.55) |
| M18 | 2181 | 16.87 (55.62) | 0.5 (1.58) |

**Table S6.** Benign CNV Regions Used for “Control” Group BD-PPI Network Comparisons. Benign CNV regions were downloaded from dbVar and used to generate BD-PPI network percentile rank distributions for “control” regions and “control” gene-sets to compare to percentile ranks for SCZ- and ASD-associated CNV BD-PPI networks.

| Chromosome | Start bp | End bp | Size (Mb) | Counts in downloaded dbVar data | Number of BrainSpan Genes | Included in Genomic-Range<sub>BD-PPI</sub> Network Analysis | Included in Gene-Set<sub>BD-PPI</sub> Network Analysis |
| --- | --- | --- | --- | --- | --- | --- | --- |
| 8 | 43541985 | 46938942 | 3.40 | 5 | 0 | 1 | 0 |
| 7 | 57927493 | 61056670 | 3.13 | 4 | 0 | 1 | 0 |
| 12 | 34854487 | 37864477 | 3.01 | 3 | 0 | 1 | 0 |
| 2 | 90321386 | 91619163 | 1.30 | 32 | 0 | 1 | 0 |
| 11 | 50121285 | 51378911 | 1.26 | 4 | 0 | 1 | 0 |
| 6 | 65259548 | 66280385 | 1.02 | 2 | 0 | 1 | 0 |
| 9 | 45721870 | 46645423 | 0.92 | 3 | 0 | 1 | 0 |
| 12 | 34089213 | 34854486 | 0.77 | 6 | 1 | 1 | 1 |
| 17 | 71847916 | 72606952 | 0.76 | 8 | 11 | 1 | 1 |
| 9 | 68759230 | 69505934 | 0.75 | 9 | 0 | 1 | 0 |
| 11 | 55213165 | 55917297 | 0.70 | 3 | 17 | 1 | 1 |
| 7 | 57230344 | 57927492 | 0.70 | 2 | 0 | 1 | 0 |
| 1 | 143719563 | 144337915 | 0.62 | 2 | 1 | 1 | 1 |
| 7 | 153074572 | 153670272 | 0.60 | 2 | 1 | 1 | 1 |
| 9 | 42456974 | 43044929 | 0.59 | 5 | 0 | 1 | 0 |
| 2 | 87376132 | 87954920 | 0.58 | 16 | 0 | 1 | 0 |
| 2 | 32677714 | 33249136 | 0.57 | 6 | 3 | 1 | 1 |
| 16 | 16292304 | 16861655 | 0.57 | 2 | 1 | 1 | 1 |
| 11 | 55980407 | 56537994 | 0.56 | 2 | 19 | 1 | 1 |
| 15 | 21381757 | 21929931 | 0.55 | 244 | 0 | 1 | 0 |
| 2 | 97714365 | 98260508 | 0.55 | 3 | 1 | 0 | 1 |
| 7 | 61307485 | 61839193 | 0.53 | 3 | 0 | 0 | 0 |
| 2 | 330817 | 861519 | 0.53 | 2 | 1 | 0 | 1 |
| 2 | 242478399 | 243007359 | 0.53 | 2 | 5 | 0 | 1 |
| 8 | 46952093 | 47457691 | 0.51 | 9 | 0 | 0 | 0 |
| 2 | 130673609 | 131172057 | 0.50 | 2 | 6 | 0 | 1 |
| 7 | 62196742 | 62690930 | 0.49 | 9 | 0 | 0 | 0 |
| 18 | 7115075 | 7608218 | 0.49 | 2 | 2 | 0 | 1 |
| 10 | 133255624 | 133728500 | 0.47 | 2 | 0 | 0 | 0 |
| 1 | 144389628 | 144860341 | 0.47 | 2 | 1 | 0 | 1 |
| 9 | 40991681 | 41461551 | 0.47 | 14 | 0 | 0 | 0 |
| 3 | 666257 | 1133628 | 0.47 | 2 | 0 | 0 | 0 |
| 8 | 42933206 | 43396776 | 0.46 | 4 | 4 | 0 | 1 |
| 17 | 21736290 | 22174114 | 0.44 | 6 | 1 | 0 | 1 |
| 6 | 168350998 | 168776932 | 0.43 | 2 | 5 | 0 | 1 |
| 12 | 33545353 | 33967851 | 0.42 | 3 | 1 | 0 | 1 |
| 19 | 23624807 | 24043127 | 0.42 | 2 | 3 | 0 | 1 |
| 22 | 16431733 | 16849852 | 0.42 | 2 | 0 | 0 | 0 |
| 2 | 83263585 | 83675297 | 0.41 | 2 | 0 | 0 | 0 |
| Y | 24238183 | 24636508 | 0.40 | 7 | 2 | 0 | 1 |

**Table S7.**
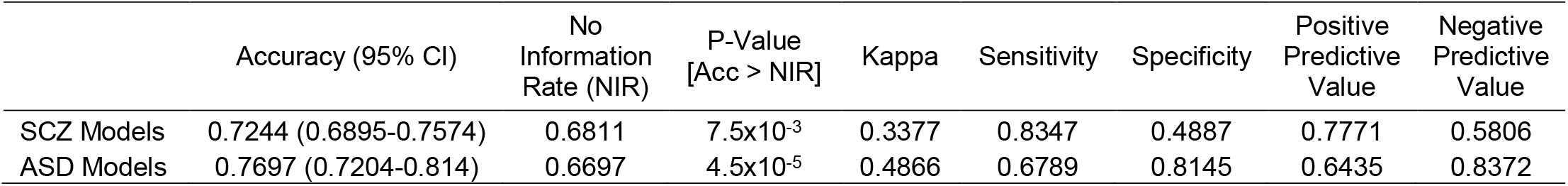
Performance of Machine Learning Models for Predicting Gene Association with SCZ and ASD. Performance statistics of linear SVM model selection process derived from nested *k*-fold cross-validation for classifying SCZ- and ASD-associated genes.

|  | Accuracy (95% CI) | No<br>Information<br>Rate (NIR) | P-Value<br>[Acc > NIR] | Kappa | Sensitivity | Specificity | Positive<br>Predictive<br>Value | Negative<br>Predictive<br>Value |
| --- | --- | --- | --- | --- | --- | --- | --- | --- |
| SCZ Models | 0.7244 (0.6895-0.7574) | 0.6811 | $7.5 \times 10^{-3}$ | 0.3377 | 0.8347 | 0.4887 | 0.7771 | 0.5806 |
| ASD Models | 0.7697 (0.7204-0.814) | 0.6697 | $4.5 \times 10^{-5}$ | 0.4866 | 0.6789 | 0.8145 | 0.6435 | 0.8372 |

**Figure S1.**
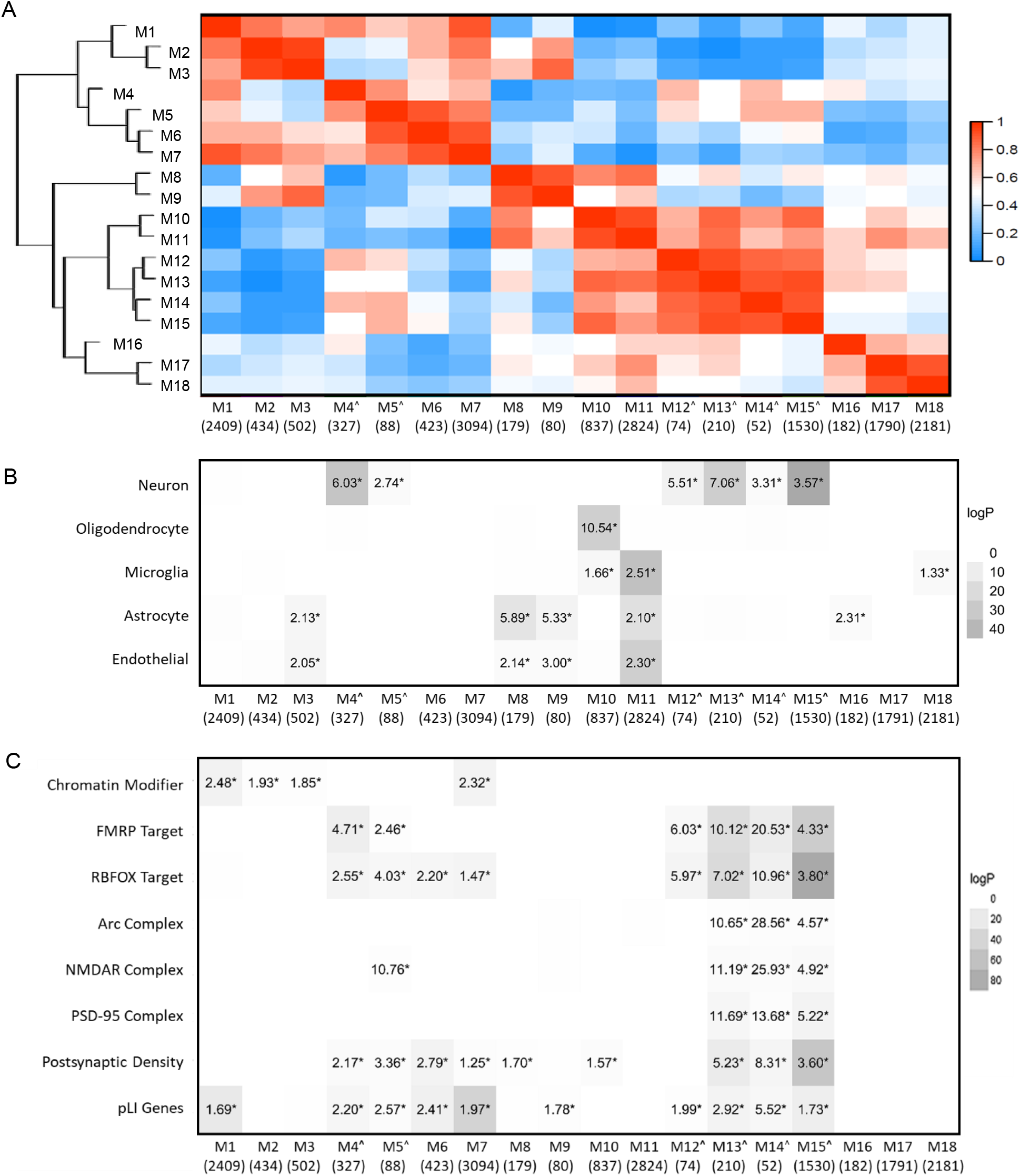
Correlation Structure of BrainSpan Modules and Enrichment for Primary Brain Cell-Types and Gene-Sets of Interest. (A) Clustering and correlation structure between 18 module eigengenes across 17116 genes derived from applying WGCNA to BrainSpan data. Modules ranged in size from 52 genes (M14) to 3094 genes (M7). Modules with high prenatal expression (M1-M7) are largely anti-correlated with modules with high postnatal expression (M10-M18). Module enrichment for (B) primary brain cell-types and (C) literature derived gene-sets of interest. Log p values are indicated by grey intensity for all enrichment odds ratios > 1.0. Numerical values for each dataset denote odds ratios for enriched modules. *FDR corrected p < 0.05 for number of modules. ^Module enriched for neuronal cell-type markers.

**Figure S2.**
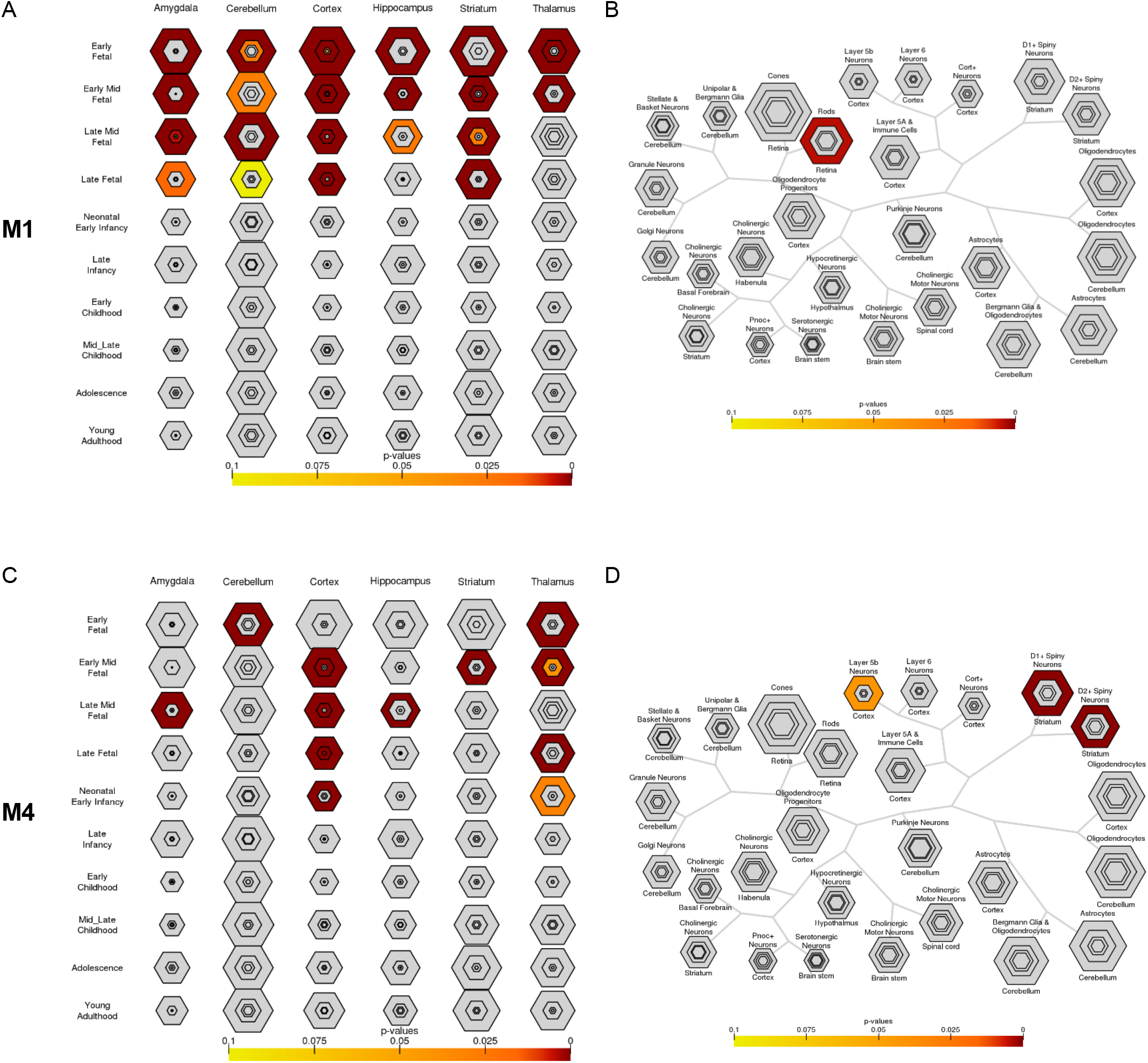
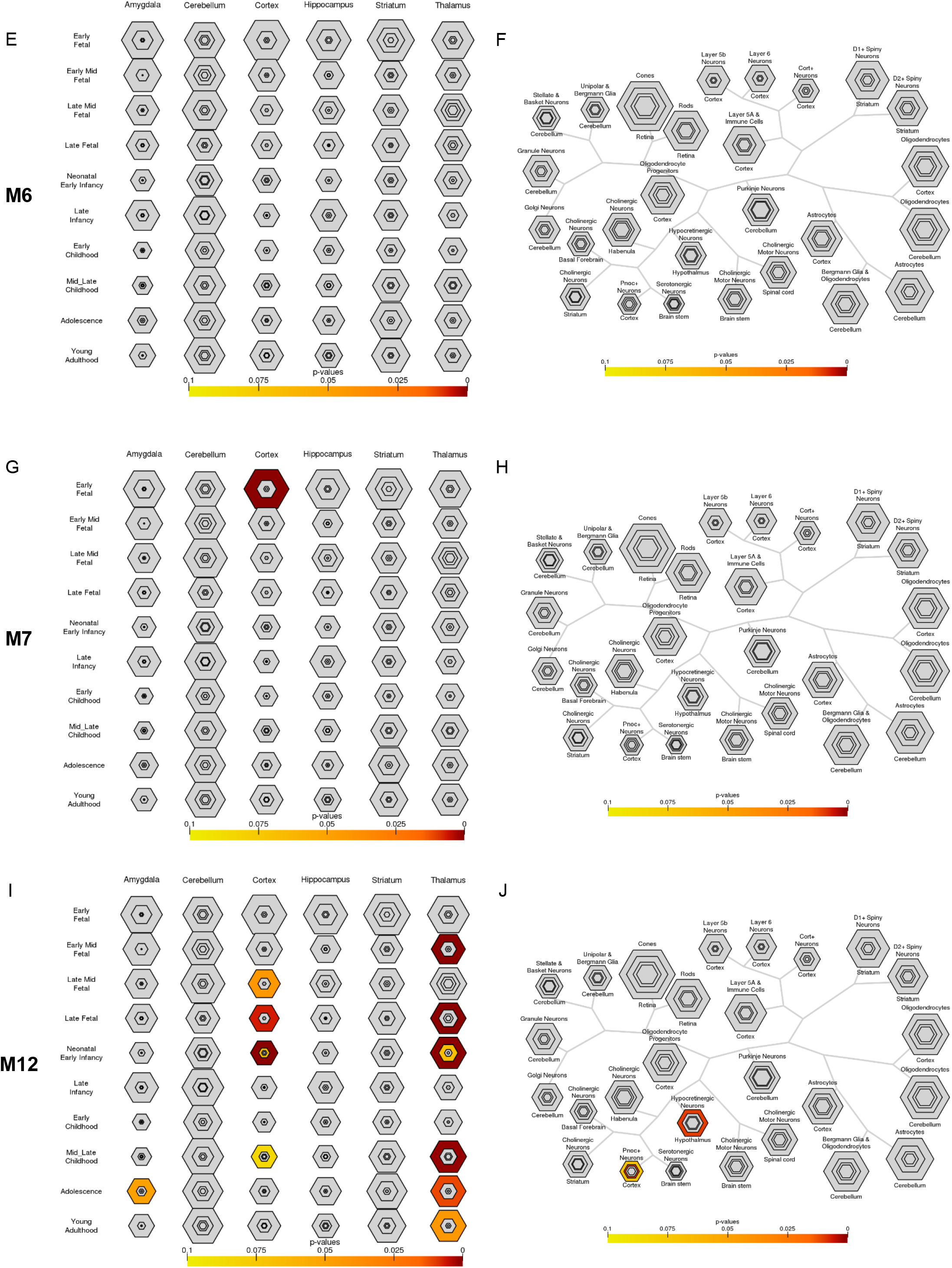
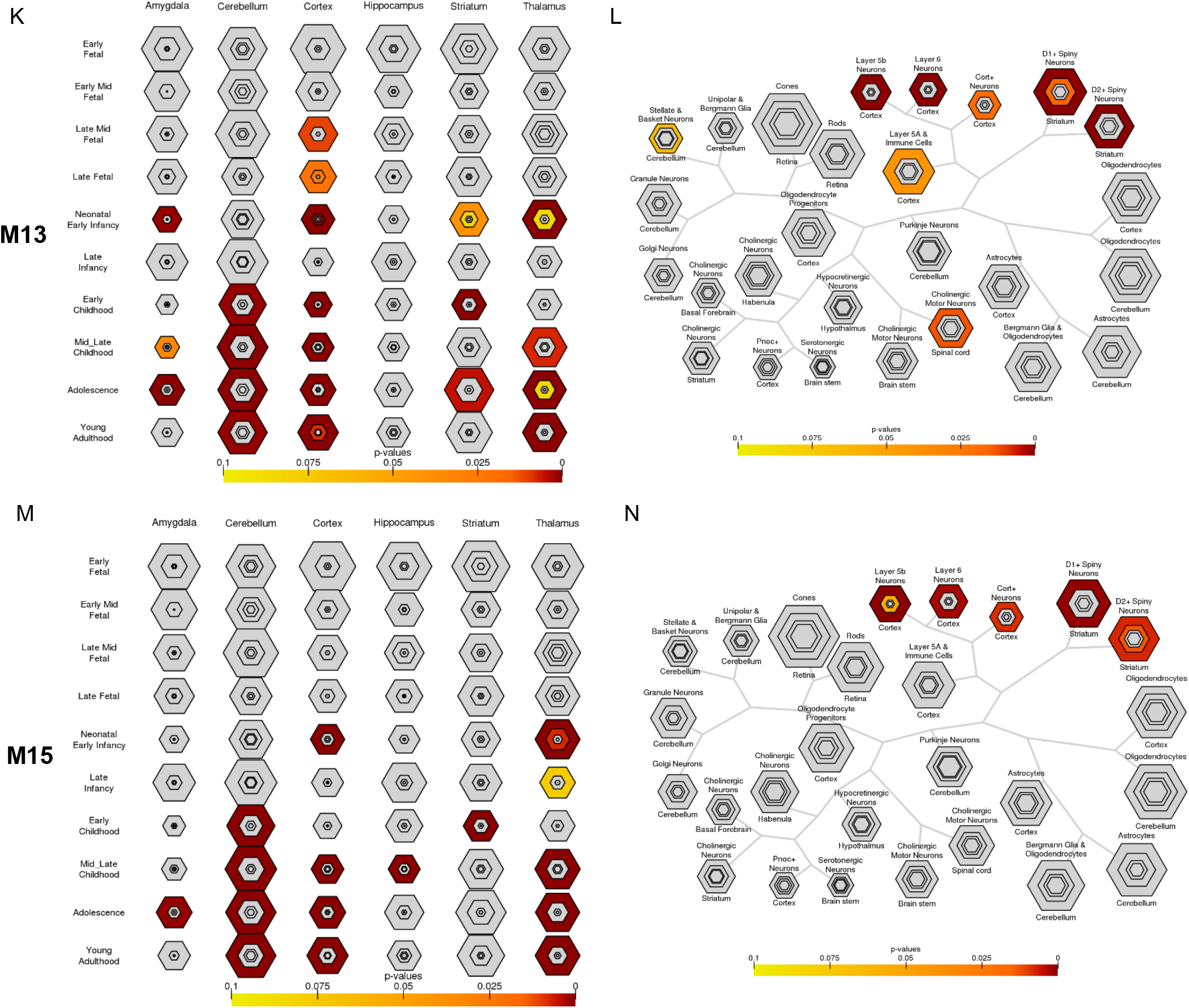
Disease-Associated BrainSpan Module Enrichment for Specific Brain Regions and Developmental Periods and for Brain Cell-Subtypes. Enrichment of the SCZ- and/or ASD-associated BrainSpan modules: (A,B) M1, (C,D) M4, (E,F) M6, (G,H) M7, (I,J) M12, (K,L) M13, and (M,N) M15 for lists of genes expressed in specific human brain regions during specific developmental periods, and in specific brain cell-subtypes, defined using varying specificity indices, was analyzed using the Specific Expression Analysis tool(13). Varying specificity thresholds are represented by the hexagon ring layers going from the least specific gene lists (outer hexagons) to the most specific gene lists (center), with hexagons scaled to the size of gene lists. BH corrected Fisher’s Exact p-values are plotted for each specificity threshold by color.

**Figure S3.**
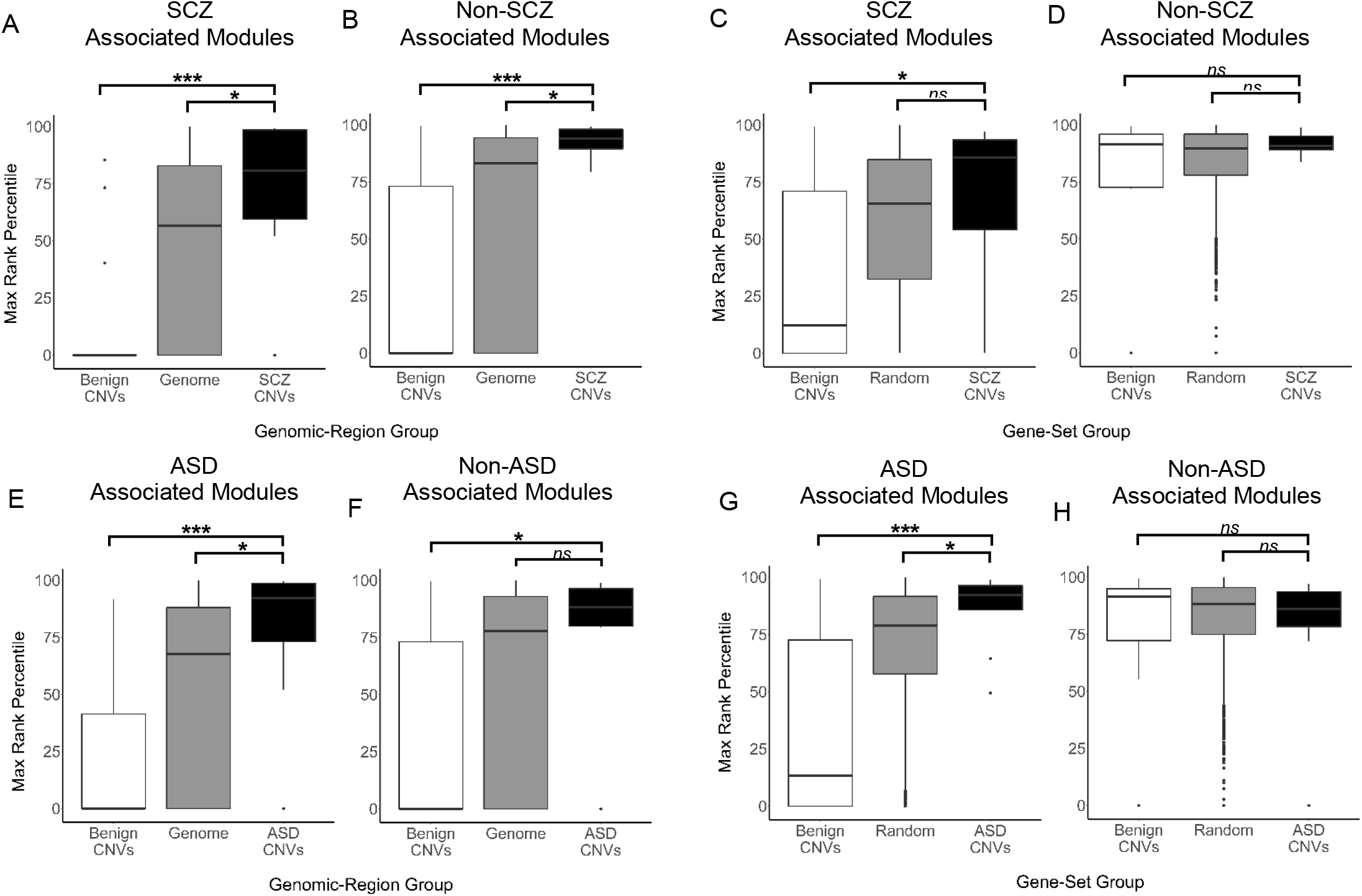
BD-PPI Networks by CNV Group for Disease and Non-Disease Associated BrainSpan Modules. Maximum percentile rank distributions of BD-PPI networks for SCZ-associated CNVs and benign CNVs, relative their comparison simulated CNVs genomic regions (i.e. equal in bp length to each SCZ-associated or benign CNV), and for all simulated CNVs generated across the genome for comparison to SCZ-associated CNVs (i.e. “Genome” Group) for: (A) SCZ- (M7, M13, M15) and (B) non-SCZ-associated modules. Maximum BD-PPI network percentile rank distributions for SCZ-associated CNVs and benign CNVs, relative to their comparison random gene-sets (i.e. equal in number of seed BrainSpan genes), and for all random gene-sets equal in number of BrainSpan genes to each SCZ-associated CNV (i.e. “Random” Group) for: (C) SCZ- and (D) non-SCZ-associated modules. Parallel BD-PPI network distributions for ASD-associated and benign CNVs, relative to their comparison equal size simulated CNVs, and for the simulated CNVs generated across the genome for comparison to ASD-associated CNVs for: (E) ASD-(M1, M4, M6, M7, M12, M13, M15) and (F) non-ASD-associated modules. Parallel distributions for BD-PPI networks for ASD-associated CNVs and benign CNVs, relative to their comparison random gene-sets equal in number of BrainSpan genes, and for all random gene-sets equal in BrainSpan genes to each ASD-associated CNV for: (G) ASD-(H) and non-ASD-associated modules. Boxplots mark the median maximum rank percentile for BD-PPI networks (when above 0) for each CNV group, with the lower and upper hinges corresponding to the 25^th^ and 75^th^ quartiles, respectively, and the lower and upper whiskers extending to 1.5 x interquartile range. Mann-Whitney U test *p < 0.05; ***p < 0.001; *ns* = non-significant.

**Figure S4.**
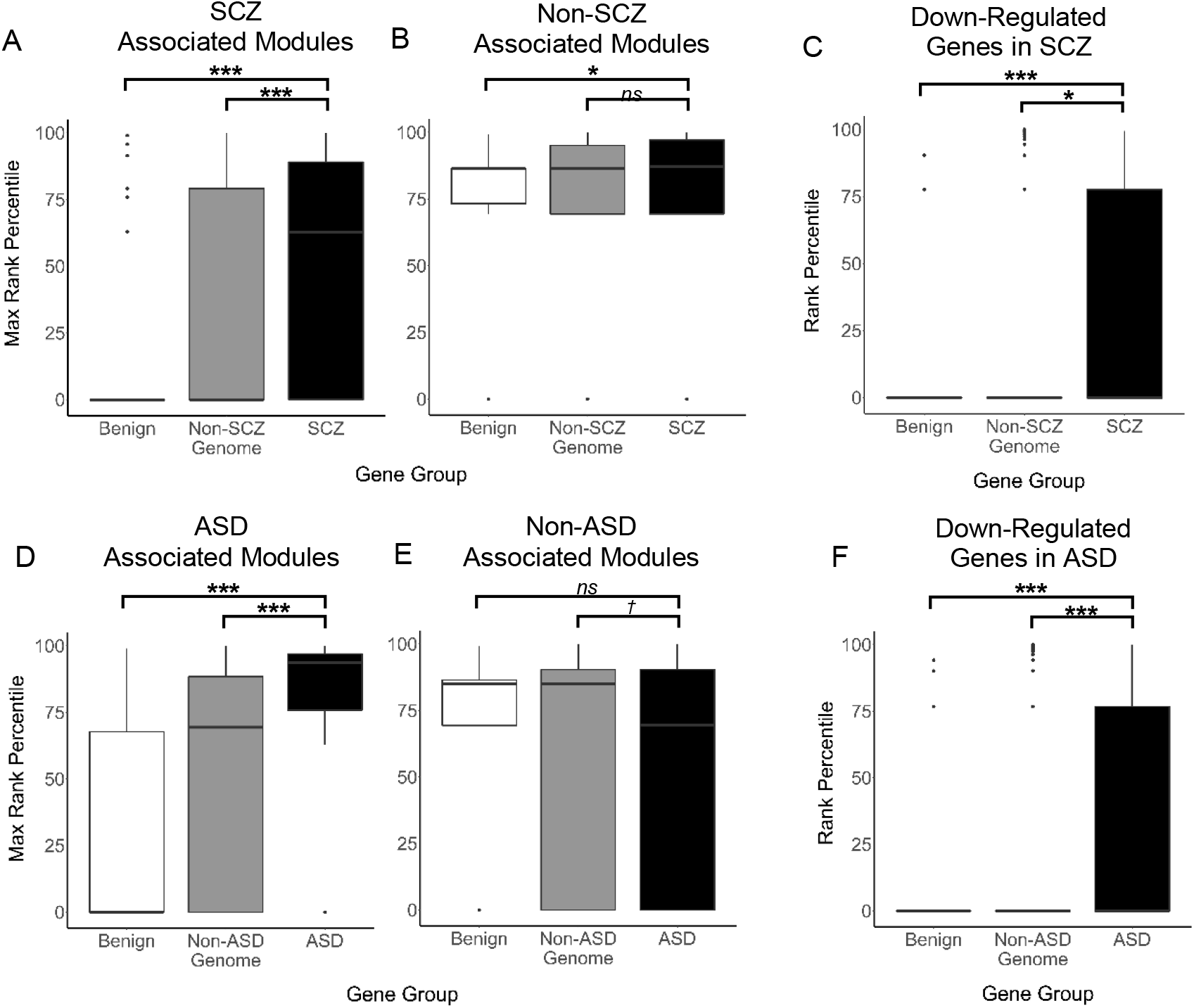
BD-PPI Networks by Gene Group for Disease and Non-Disease Associated BrainSpan Modules and Key Pathophysiological Signatures. Maximum percentile rank distributions of BD-PPI networks for SCZ-associated genes versus non-SCZ-associated genes and “control” genes in benign CNVs across (A) SCZ-associated modules (M7, M13, M15) and (B) non-SCZ-associated modules (all other modules); and overlapping (C) down-regulated genes in SCZ cortex. Parallel plots for ASD-associated genes versus non-ASD-associated genes and “control” genes in benign CNVs across (D) ASD-associated modules (M1, M4, M6, M7, M12, M13, M15); (E) non-ASD-associated modules (all other modules); and overlap with (F) down-regulated genes in ASD cortex. Boxplots mark the median maximum rank percentile for BD-PPI networks (when above 0), with the lower and upper hinges corresponding to the 25^th^ and 75^th^ quartiles, respectively, and the lower and upper whiskers extending to 1.5 x interquartile range. SCZ or ASD group higher than comparison group *p < 0.05; ***p < 0.001; *ns* = non-significant. Comparison group higher than SCZ or ASD group ^†^p< 0.05.

**Figure S5.**
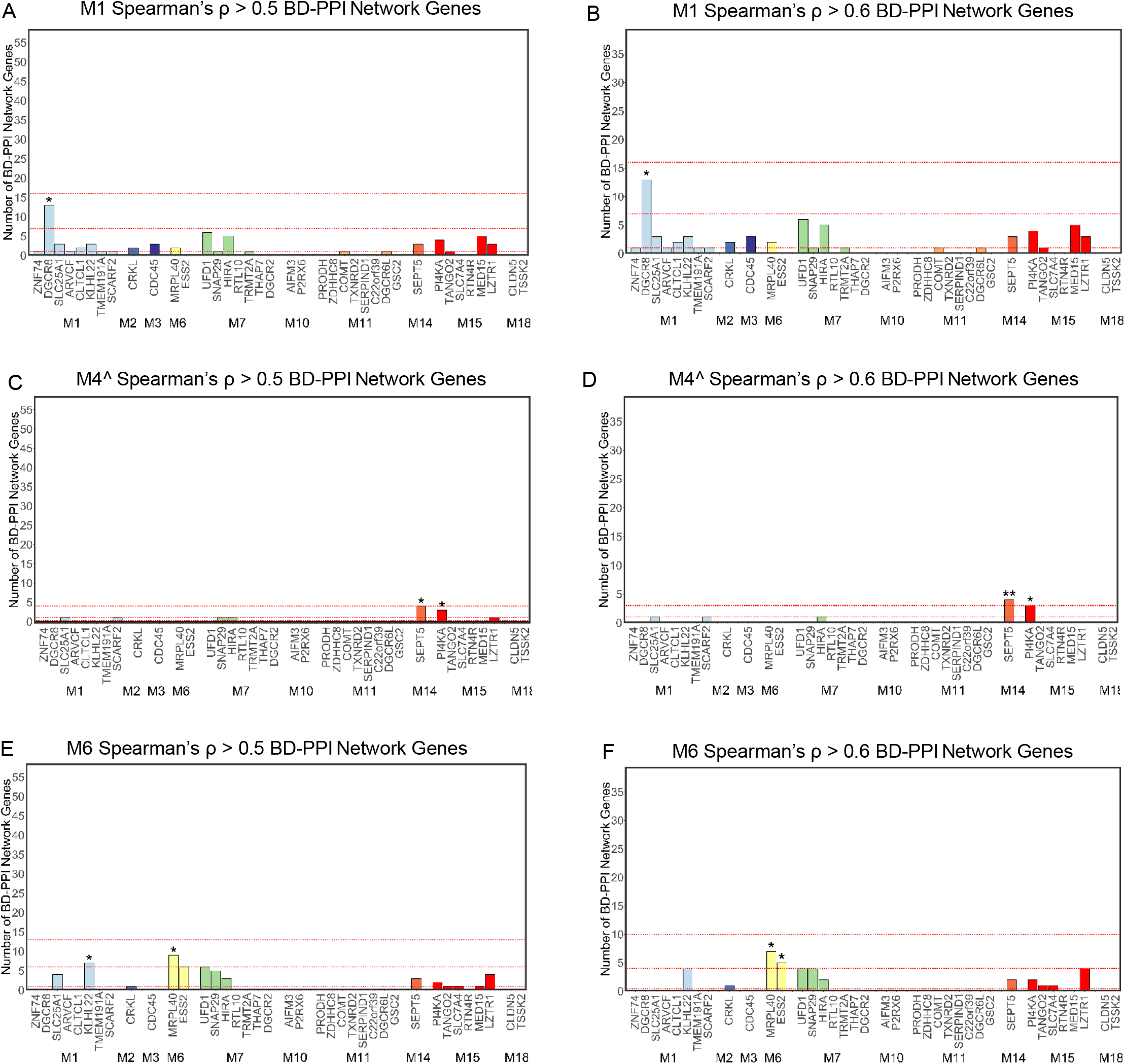
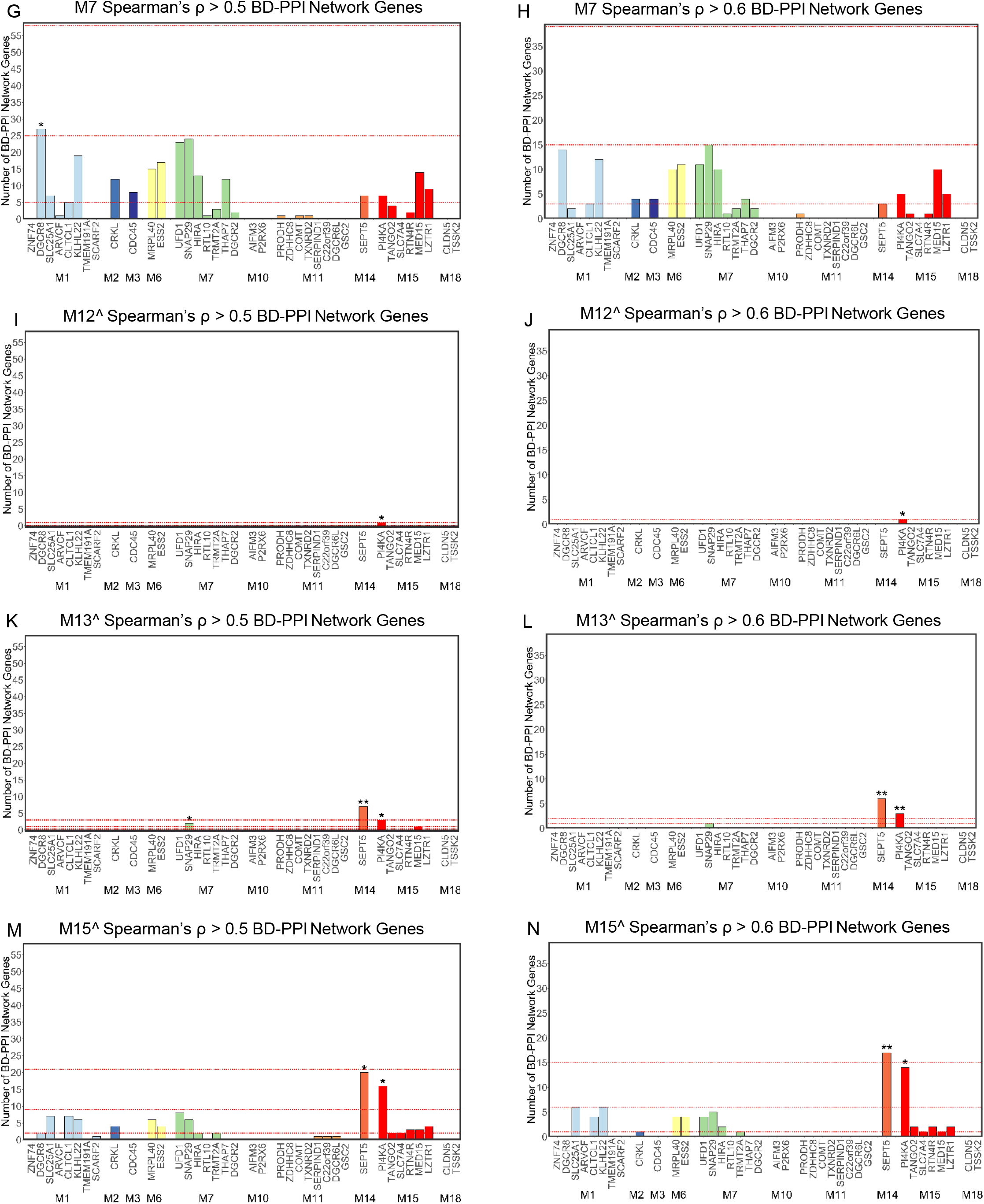
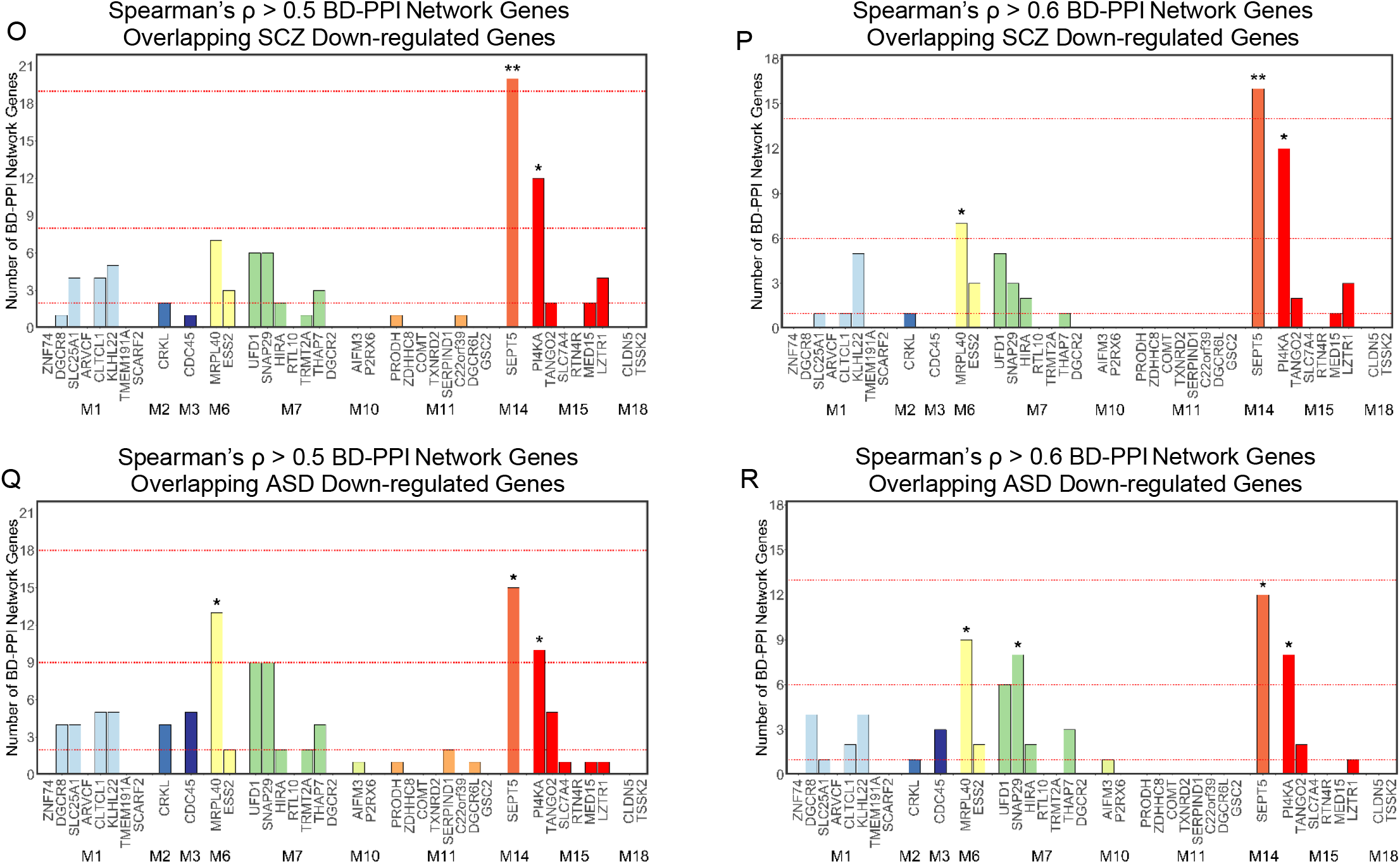
Robustness of Prioritized 22q11.2 Genes Across Varying BD-PPI Network Definitions. Number of BD-PPI network genes for each 22q11.2 gene, defined using varying Spearman’s ρ thresholds, for the SCZ- and/or ASD-associated modules: M1 (A,B), M4 (C,D), M6 (E,F), M7 (G,H), M12 (I,J), M13 (K,L), and M15 (M,N), and overlapping significantly down-regulated genes in (O,P) SCZ and (Q,R) ASD post-mortem cortex. In each plot, the 22q11.2 genes are shown grouped by module membership. The top, second, and third red dotted lines on each plot mark the number of genes nearest to the 99^th^, 95^th^, and 75^th^ percentiles, respectively, for BrainSpan genes, when above 0. Genes above the 95^th^ or 99^th^ percentile for each module and gene expression signature are denoted by * or **, respectively. ^Module enriched for neuronal cell-type markers.

**Figure S6.**
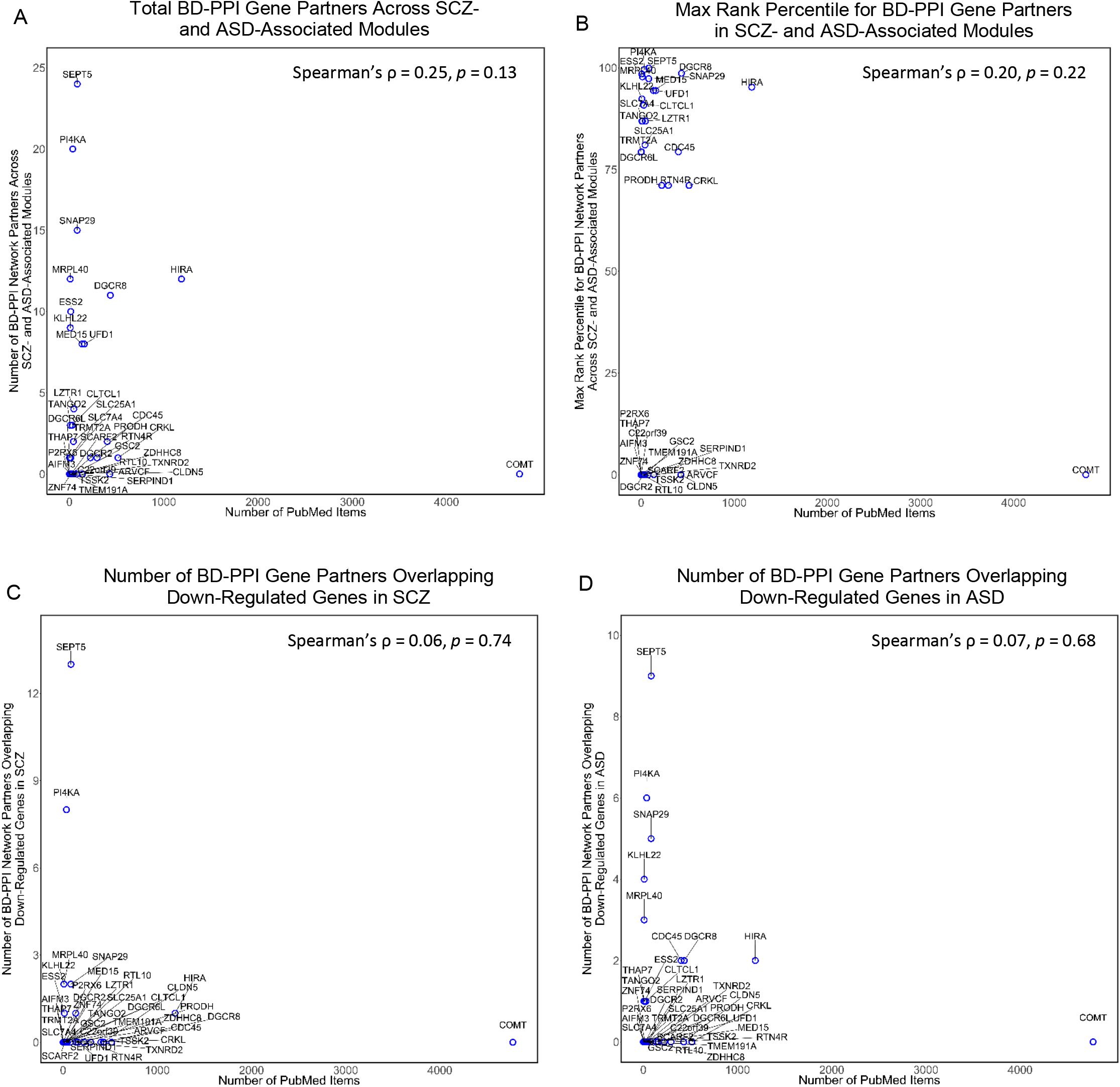
Study Bias Does Not Account for Prioritization of 22q11.2 Genes. Spearman’s rank correlations between number of PubMed entries per 22q11.2 gene and (A) total number of BD-PPI network gene partners across all SCZ- and ASD-associated modules (M1, M4, M6, M7, M12, M13, M15); (B) maximum rank percentile for BD-PPI network partners in any SCZ- or ASD-associated module; (c) number of BD-PPI network partners overlapping significantly down-regulated genes in SCZ; and (D) number of BD-PPI network partners overlapping significantly down-regulated genes in ASD. There were no significant relationships between number of PubMed entries for each 22q11.2 gene and any disease-related metric of interest.

